# Antiviral DExD/H-box helicase 60 selectively inhibits translation from type II internal ribosome entry sites

**DOI:** 10.1101/2022.02.08.479557

**Authors:** Mohammad Sadic, William M. Schneider, Olga Katsara, Gisselle N. Medina, Aishwarya Mogulothu, Yingpu Yu, Meigang Gu, Teresa de los Santos, Robert J. Schneider, Meike Dittmann

**Author notes:** Competing interests: The authors declare that no competing interests exist.

## Abstract

During viral infection, competition ensues between viruses and their host cells to control the protein synthesis machinery. In response, certain host defense proteins globally limit mRNA translation. However, this is also detrimental for host protein synthesis. Here we describe an interferon-stimulated helicase, DDX60, that specifically inhibits translation from type II viral internal ribosome entry sites (IRESs). IRESs are RNA structures that enable mRNAs to recruit ribosomes directly, bypassing translation initiation using a 5’ cap. DDX60 was previously observed to inhibit replication of a reporter hepatitis C virus (HCV). We show that DDX60 likely does not inhibit HCV replication, but surprisingly, inhibits the type II IRES used in the reporter HCV genomic RNA. Using firefly luciferase mRNAs translationally driven by different viral IRESs or a 5’ cap analog, we show that DDX60 selectively reduces translation driven by type II IRESs of encephalomyocarditis virus (EMCV) and foot and mouth disease virus (FMDV), but not other IRES types or a 5’ cap analog. Correspondingly, DDX60 reduces EMCV and FMDV (type II IRES) replication, but not poliovirus or bovine enterovirus 1 (type I IRES) replication. Furthermore, replacing the IRES of poliovirus with a type II IRES is sufficient for DDX60 to inhibit poliovirus replication. Finally, we demonstrate that DDX60 specifically reduces polysome binding on type II IRES mRNA, but not 5’ cap-dependent mRNA. Our data demonstrate that the cellular defense factor DDX60 counteracts viral takeover of host translation by blocking ribosome access to type II IRES elements specifically.

## Introduction

Co-opting host cell protein synthesis is a hallmark of many virus infections. In eukaryotes, a covalent m^7^GpppG 5’ cap structure on host messenger RNAs (mRNAs) enables the recruitment of a translation initiation factor complex that recruits the 40S ribosome subunit to initiate mRNA translation (Jackson et al., 2010; Merrick & Pavitt, 2018). The cap structure is initially recognized by a eukaryotic initiation factor (eIF) protein eIF4E, which forms a complex with the scaffold protein eIF4G. An interaction between eIF4G and eIF3 then assembles a 43S preinitiation complex consisting of a 40S ribosomal subunit bound to eIF3, eIF1, eIF1A, and a ternary complex of GTP bound eIF2 and initiator Met-tRNA_*i*_^Met^, among other factors. This ribosomal complex then scans the mRNA in the 5’ to 3’ direction. During scanning, eIF4G-bound RNA helicase eIF4A and activator protein eIF4B unwind RNA secondary structures in the mRNA until the start codon is identified. Subsequently eIF1, eIF1A, and eIF5 assist in positioning the 40S ribosomal subunit such that the initiator Met-tRNA_*i*_^Met^ is at the peptidyl (P)-site of the 40S ribosomal subunit. eIF5 then promotes GTP hydrolysis by eIF2, releasing eIF2 and eIF5 for subsequent cycles of translation initiation. Lastly, the GTPase eIF5B assists in joining the 60S ribosomal subunit to the 40S subunit to form an 80S initiation complex. The poly A-binding protein (PABP) interacts with the 3’-poly(A) tail and eIF4G, further promoting mRNA translation initiation.

Viruses have developed diverse mechanisms to compete with and dominate the host protein synthesis machinery, much of it centered on maintaining cap-dependent mRNA translation or bypassing it completely. Some viruses utilize eukaryotic capping enzymes to add a m^7^Gppp 5’ cap to their mRNAs, while others encode their own viral capping enzymes to add a 5’ cap that functionally mimics a eukaryotic 5’ cap. Some viruses naturally have uncapped mRNAs but can “snatch” capped 5’ terminal fragments from host mRNAs (Decroly et al., 2012; Plotch et al., 1981), while others covalently link their uncapped mRNA to a 5’ terminal protein that mechanistically acts like a 5’ cap to recruit translation initiation complex proteins (Goodfellow et al., 2005). Other viruses directly recruit ribosomes to the mRNA and bypass the requirement for 5’ cap recognition using structured RNA elements called IRESs (Jang et al., 1988; Pelletier & Sonenberg, 1988; Stern-Ginossar et al., 2019). IRESs assemble the translation initiation apparatus either upstream of or at an initiation codon, independently of a 5’ cap structure (Fraser & Doudna, 2007; Lee et al., 2017; Lozano & Martínez-Salas, 2015; Martinez-Salas et al., 2018; Yamamoto et al., 2017).

IRESs assemble the translation initiation apparatus by interacting with a defined set of eIFs and host IRES transacting factor proteins (ITAFs) that assist in the recruitment of the 40S ribosomal subunit. Viral IRESs are classically divided into four subtypes based on their unique RNA structures, differential requirements for eIFs and ITAFs, and start codon recognition mechanisms (Beales et al., 2003; Belsham, 1992; Hunt et al., 1993; Kaminski et al., 1990; Lee et al., 2017; Lozano & Martínez-Salas, 2015; Martinez-Salas et al., 2018; Ohlmann & Jackson, 1999; Yamamoto et al., 2017). Viral type I IRESs found in picornaviruses such as poliovirus and enterovirus 71 (EV71), for example, employ a ribosomal scanning mechanism for start codon recognition with the assistance of eIFs 1A, 2, 3, 4A, 4B, central domain of 4G, and ITAFs PCBP1/2, PTB, hnRNPA1, and other proteins (Martinez-Salas et al., 2018; Pelletier & Sonenberg, 1988; Stern-Ginossar et al., 2019; Thompson & Sarnow, 2003). Type II IRESs found in encephalomyocarditis virus (EMCV) and foot and mouth disease virus (FMDV) on the other hand direct ribosome entry at an AUG in the 3’ end of the IRES, or one located a short distance away with the assistance of the same set of eIFs as type I IRESs, but a different set of ITAFs (Belsham, 1992; Jang et al., 1988; Martinez-Salas et al., 2018; Stern-Ginossar et al., 2019). Type III IRESs found in some flaviviruses such as HCV and bovine viral diarrhea virus (BVDV) recruit the 40S ribosomal subunit close to the start codon without the use of eIFs, and subsequently recruit GTP bound eIF2 and initiator Met-tRNA_*i*_^Met^ and eIF3 to facilitate 60S ribosomal subunit joining (Fraser & Doudna, 2007; Sanderbrand et al., 2000; Tsukiyama-Kohara et al., 1992). Finally, type IV IRESs found in the dicistrovirus cricket paralysis virus (CrPV), require no eIFs or ITAFs for 40S ribosomal subunit recruitment, and initiate translation at a noncanonical start codon from the A-site of the ribosome (Jan & Sarnow, 2002; Wilson et al., 2000). Cap independent translation mechanisms used by IRESs are well suited to initiate protein synthesis even under conditions of cellular stress when eIF4E-binding protein (4E-BP) family members bind and sequester eIF4E to prevent its binding to eIF4G (Haghighat et al., 1995; Stern-Ginossar et al., 2019). Many viruses with IRESs also developed mechanisms to selectively favor translation of their own mRNAs over their host’s cap-dependent mRNAs (Belsham & Brangwyn, 1990; Garaigorta & Chisari, 2009; Gradi et al., 1998; Jan et al., 2016; Mohr & Sonenberg, 2012; Stern-Ginossar et al., 2019).

While viruses must often compete for the host’s translation machinery, cells respond by enacting different mechanisms to block overall protein synthesis, and in some cases specifically inhibit translation of viral mRNAs. Interferons (IFNs), produced by cells upon viral infection, trigger the expression of a variety of interferon stimulated genes (ISGs) that have diverse antiviral functions, some of which target translation (Diamond & Farzan, 2013; Ficarelli et al., 2021; Hoffmann et al., 2015; Hopfner & Hornung, 2020; Li & Wu, 2021; Schneider et al., 2014; Schoggins, 2019; Stern-Ginossar et al., 2019). Among them is the double-stranded RNA activated protein kinase PKR, which phosphorylates the eIF2 α-subunit to impair GDP to GTP exchange by the eIF2B GTP exchange factor, thus inhibiting global protein synthesis. Another mechanism involves activation of oligoadenylate synthase (OAS), which synthesizes short oligoadenylate polymers to stimulate RNase L to indiscriminately degrade ribosomal and both viral and some host mRNAs (Burke et al., 2019). The interferon-induced protein with tetratricopeptide repeats (IFIT) family members and interferon-induced transmembrane protein (IFITM) family members bind specific eIFs to restrict global protein synthesis or recognize structures absent in viral 5’ caps such as 2’O-methylation. Finally, the zinc finger antiviral protein (ZAP) triggers viral RNA degradation and limits interactions between certain eIFs. While these mechanisms limit translation of viral mRNAs, they come at the cost of downregulating host protein synthesis. Here, we describe the ISG DExD/H-box helicase 60 (DDX60), an RNA helicase that can inhibit viral type II IRES-driven translation while leaving host 5’ cap-driven mRNA translation intact.

The antiviral function of DDX60 was initially discovered in a screen for antiviral ISGs, where it was shown to inhibit a reporter HCV (Schoggins et al., 2011). Later studies probing for the antiviral mechanism of DDX60 generated conflicting data. One group found DDX60 to act as a sentinel for the viral RNA recognition receptor retinoic acid-inducible gene-I (RIG-I) (Miyashita et al., 2011), and to promote degradation of viral RNA independently of RIG-I (Oshiumi et al., 2015). However, another group presented evidence against a role for DDX60 as a sentinel for RIG-I (Goubau et al., 2015), suggesting instead that DDX60 may enact a specific antiviral mechanism for one or a small group of viruses.

Here, we aimed to clarify the mechanism for DDX60 antiviral activity. We first show that upon IFN-ß treatment, DDX60 has prolonged and delayed expression dynamics at the mRNA and protein levels, respectively. Through mutagenesis and antiviral assays, we demonstrate that N- and C-terminal regions and predicted helicase and ATP binding motifs in DDX60 are important for its antiviral activity. We next use comparative antiviral experiments to show that DDX60 targets type II IRESs found in a group of viruses. We generated *in vitro* transcribed mRNA reporters to demonstrate that DDX60 specifically inhibits the type II family of IRESs and further show that the type II IRES is sufficient to confer virus inhibition by DDX60. Lastly, we found that DDX60 inhibits type II IRES containing mRNAs, not by affecting the mRNA abundance, but by reducing polysome binding on these mRNAs. Importantly, DDX60 shows no effect on the overall translation status of the cell nor an effect on the translation of *in vitro* synthesized 5’ capped mRNA. Our work suggests that DDX60 acts as an ISG that inhibits type II IRES-mediated mRNA translation and can discriminate between 5’ cap-independent and dependent translation mechanisms. Studying the anti-IRES mechanism of DDX60 could lead to novel strategies for targeting certain virus translation mechanisms.

## Results

### DDX60 displays dynamics of a type I ISG at the mRNA and protein level in multiple cell lines

DDX60 is part of a large program of genes activated by IFN to confer protection against virus infection. Indeed, gene expression of *DDX60* at the mRNA level is triggered by various stimuli in human cell lines and mouse tissue, including poly(I:C), type I IFN, and virus infections (Goubau et al., 2015; Miyashita et al., 2011). To recapitulate the role of DDX60 as a de facto ISG, we analyzed the dynamics of *DDX60* mRNA in four different human cell lines upon treatment with IFN-ß. We treated three different epithelial cell lines – HEK293T (human embryonic kidney), A549 (lung adenocarcinoma), and HeLa (cervical adenocarcinoma) – and primary fibroblasts –HFF1 (human foreskin fibroblast) – with IFN-ß for either 0, 6, 12, 24, or 48 hours and subsequently analyzed *DDX60* expression using RT-qPCR. *Interferon regulatory factor 1* (*IRF1*), an IFN stimulated transcription factor with broad antiviral function and known expression dynamics (Feng et al., 2021; Forero et al., 2019; Schoggins et al., 2011), served as a control. In all cell lines tested, *IRF1* acted as observed previously, with mRNA expression peaking at 6 to 12 hours post IFN-ß stimulation, and then decreasing back to baseline levels after 12 hours (**Fig. S1, line graphs on the right**). *DDX60* mRNA levels peaked to even higher levels, on average, than *IRF1* and showed varied time kinetics at the mRNA level in the different cell lines (**Fig. S1A-D**). In HFF1, *DDX60* mRNA dynamics resembled that of IRF1, peaking at 12 hours post-IFN-ß treatment before returning to baseline levels (**Fig. S1C, left line graph**). In HEK293T, A549, and HeLa cells, however, *DDX60* expression peaked later at 12-24 hours post-IFN-ß treatment and decreased steadily, never fully reaching baseline levels even after 48 hours post-IFN-ß treatment (**Fig. S1A, B, and D left line graphs**). Unlike primary HFF1, HEK293T, A549, and HeLa cells are cancer cell lines with varying chromosome copy numbers, active signaling pathways, and dysregulated transcriptional and post-transcriptional mechanisms (Lopes-Ramos et al., 2017; Pan et al., 2009). This could in part explain the observed *DDX60* expression differences in cancer cell lines compared to a primary fibroblast line. Overall, our IFN stimulation and mRNA analysis demonstrate that *DDX60* displays general characteristics of an ISG.

We next sought to determine if protein levels of endogenous DDX60 also change with IFN treatment. We thus analyzed DDX60 protein dynamics in our four IFN-ß treated cell lines by western blot. Expectedly, DDX60 protein production increased approximately 3 to 10-fold from a baseline of little to no expression with IFN-ß treatment (**Fig. S1E-H**). However, compared to *DDX60* mRNA levels, DDX60 protein levels showed delayed expression dynamics in all cell lines, peaking at 24 or even 48 hours post-IFN-ß treatment (**Fig. S1E-H**).

Together, our findings show that DDX60 is an ISG with very low to undetectable concentrations at baseline that then peak at both the RNA and protein levels after IFN treatment in various human cell culture systems.

**Figure S1.**
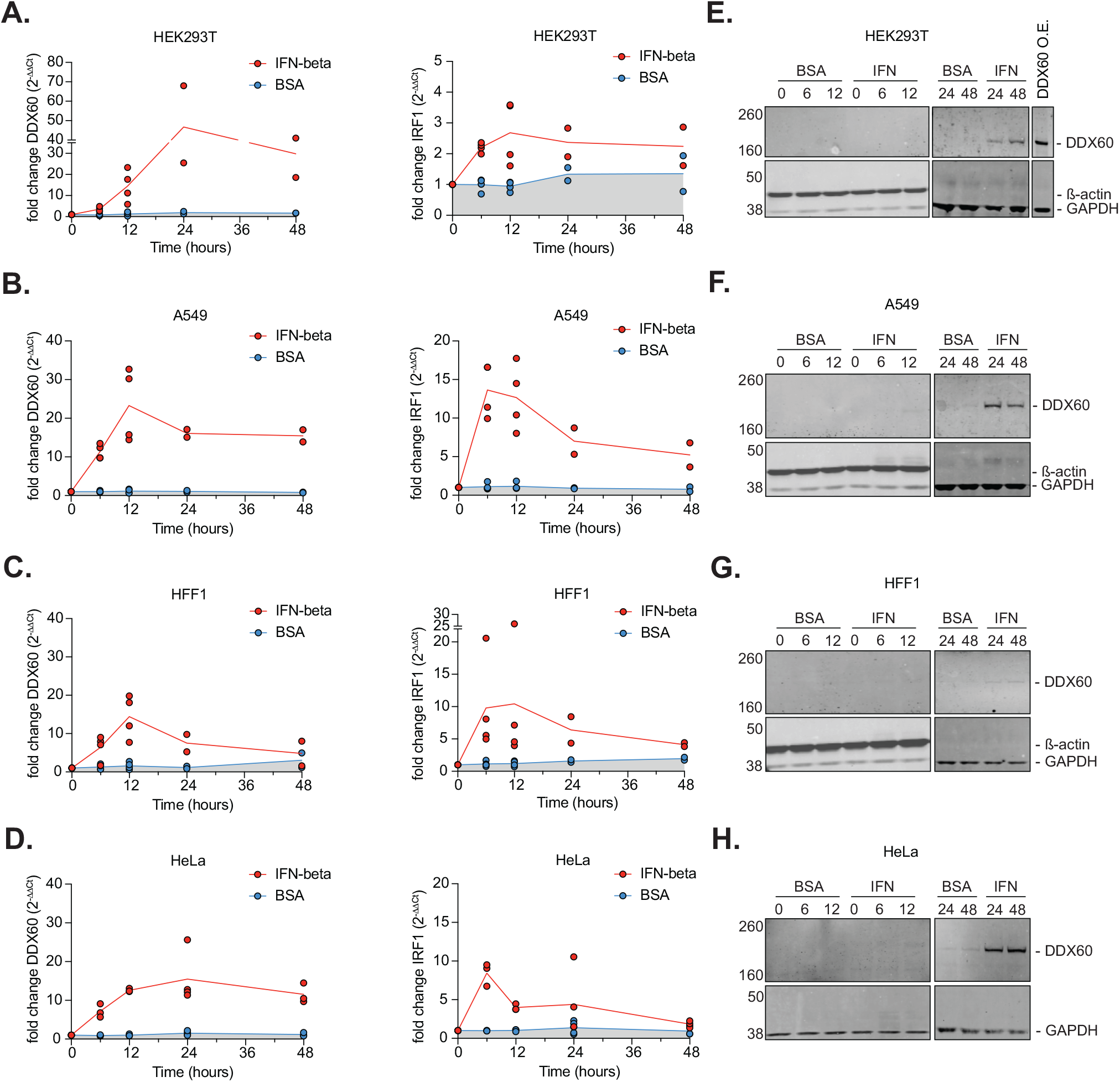
DDX60 mRNA and protein expression upon interferon treatment. **(A, E)** HEK293T, **(B, F)** A549, **(C, G)** HFF1, or **(D, H)** HeLa cells treated with either 0.1% BSA (carrier control) or 500 U/mL of interferon-ß for 0, 6, 12, 24, or 48 hours. Cells were harvested for mRNA analysis (A – D) using RT-qPCR or protein analysis (E–H) using western blot. Panel E includes one HEK293T sample transfected with DDX60 wild type run on the same gel as 24- and 48-hour time points to show relative DDX60 protein levels in interferon-ß treated versus transfected cells. All western blots for 0-, 6-, and 12-hour time points were run on the same gel but are separated by cell line for visualization purposes. All western blots for the 24- and 48-hour time points were run on the same gel but are separated by cell line for visualization purposes. Data is representative of at least 2 biological replicates per time point from experiments performed on different days.

### DDX60 inhibits replication of a bicistronic reporter HCV carrying an EMCV IRES

Previous studies showed that DDX60 inhibits replication of a bicistronic reporter HCV (Oshiumi et al., 2015; Schoggins et al., 2011). To begin determining how DDX60 does so, we used InterPro and published literature (Johnson & Jackson, 2013; Pause & Sonenberg, 1992; Pyle, 2008; Schwer & Meszaros, 2000; Umate et al., 2011), to identify potential functional domains and motifs (**Fig. 1A**). We then introduced N-terminal and C-terminal truncations and single point mutations in residues predicted to confer ATP binding/hydrolysis and helicase activity to DDX60. We first confirmed that all mutants could be ectopically expressed to equal levels by western blot (**Fig. 1B**). We then used a previously developed virus inhibition assay to assess the antiviral capacity of the different DDX60 constructs (Schoggins et al., 2011). Briefly, we transfected Huh-7 cells with wild type or mutant DDX60 plasmid containing a red fluorescent protein (RFP) marker to monitor transfection efficiency. We then challenged transfected cells with a yellow fluorescent protein (Ypet)-expressing reporter HCV at a dose yielding approximately 50% infected (Ypet+) cells in our negative control, as previously determined by flow cytometry-based infectivity assays (Jones et al., 2010; Schoggins et al., 2011). The percentage of Ypet-positive cells (infected) within the RFP-positive population (transfected) at 72 h post infection was assayed by flow cytometry. Firefly luciferase (Fluc) served as a negative control and normalization factor and the broad transcriptional activator of antiviral genes, IRF1 (Feng et al., 2021), served as a positive control for virus inhibition. As previously observed (Schoggins et al., 2011), IRF1, a potent antiviral protein, reduced the percentage of infected (Ypet+) cells by approximately 80% and wild type DDX60 reduced the percentage of infected cells by approximately 30% relative to our Fluc control. Furthermore, we found that predicted ATP binding residues, helicase motif, and N and C-terminal extensions are all important for DDX60’s antiviral activity (**Fig. 1C**), suggesting that ATP binding/hydrolysis, RNA unwinding, and the highly conserved DExD/H motif may be important for virus inhibition by DDX60. Equally important are functions associated with N and C-terminal extensions, which in other RNA helicases, allow for protein-protein interactions during RNA substrate recognition (Lingaraju et al., 2019; Thoms et al., 2015; Wang et al., 2005).

**Figure 1.**
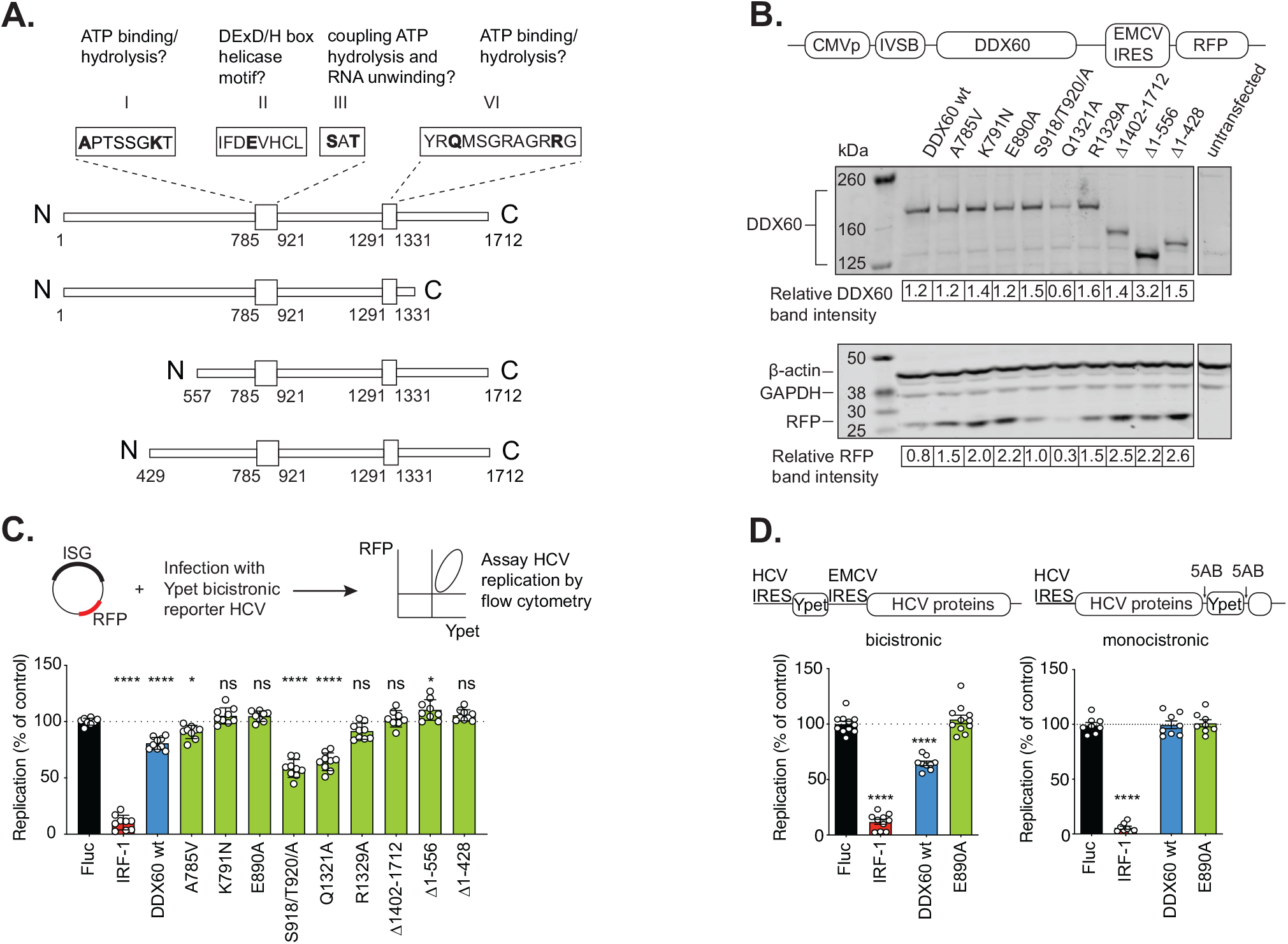
Functional mapping of DDX60 antiviral domains and interrogation of anti-HCV activity. **(A)** Schematic of DDX60 protein with putative functional domains. Helicase ATP binding type I domain (amino acids 785 - 921) and C-terminal helicase domain (amino acids 1291-1331) are shown as larger boxes in linear DDX60 schematic. Amino acids are numbered below. Putative functional motifs (I, II, III, and VI) are annotated. The function of amino acids in bold as well as N and C-terminal regions were interrogated in antiviral assays. (**B**) Assessment of exogenous DDX60 expression. HEK293T cells transfected with DDX60 wild type (wt), or DDX60 mutants and analyzed by western blot for DDX60, ß-actin and GAPDH (loading controls), and RFP (reporter). DDX60 and RFP quantification by densitometry are shown below. **(C)** HCV antiviral assays with DDX60 wt or mutant panel. Huh-7 cells transfected with an RFP containing plasmid backbone encoding either Firefly luciferase (Fluc, negative, and transfection control), IRF1 (positive antiviral control), DDX60 wt, or DDX60 mutants and challenged with HCV-Ypet, a bicistronic reporter HCV where Ypet reporter protein is driven by HCV IRES and HCV polyprotein consisting of C, E1, E2, p7, NS2, NS3, 4A, 4B, NS5A, and NS5B is driven by EMCV IRES. **(D)** Effect of DDX60 on replication of bicistronic or monocistronic reporter HCVs. Huh-7 cells transfected as in (C) and infected with either bicistronic HCV-Ypet (left) or monocistronic HCV J6/JFH-5AB-YPet. Ypet reporter in monocistronic HCV is placed in between NS5A and NS5B. For (C) and (D), percent of Ypet+ cells in RFP+ cells is scaled to one replicate of Fluc control. Data shows mean ± SD for at least 3 biological replicates from experiments performed on separate days; ns = not significant, *p < 0.05, ****p < 0.0001 using ANOVA and Dunnett’s multiple comparison test against Fluc.

For these initial experiments, we used the same reporter HCV as in a screen for ISGs that initially identified DDX60 to be antiviral (Schoggins et al., 2011). This reporter HCV is bicistronic, as translation of the Ypet reporter is driven by the HCV internal ribosome entry site (IRES) and translation of the HCV polyprotein is subsequently driven by an inserted encephalomyocarditis virus internal ribosome entry site (EMCV IRES) (**Fig. 1D** (Jones et al., 2010; Jones et al., 2007; Schoggins et al., 2011). To validate our findings and rule out artifactual observations due to the use of a reporter virus encoding a foreign viral element, we employed our flow cytometry-based virus inhibition assay using a monocistronic reporter HCV where translation is initiated by the endogenous HCV IRES and the Ypet is translated as a part of the HCV polyprotein and subsequently excised due to flanking NS5AB cleavage sites (**Fig. 1D** (Horwitz et al., 2013; Jones et al., 2007). While DDX60 successfully inhibited replication of the bicistronic reporter HCV as observed previously (**Fig. 1C, Fig. 1D, left panel**, (Schoggins et al., 2011), DDX60 failed to inhibit the monocistronic reporter HCV (**Fig. 1D, right**). Additionally, the EMCV IRES-driven RFP encoded in our plasmid constructs used in Figure 1B and 1C showed expression levels that mimicked the expression of our bicistronic Ypet reporter HCV, but 5’ cap-driven proteins like ß-actin, GAPDH, or our DDX60 transgenes of interest did not (**Fig. 1B**). As the main distinguishing feature between the two reporter HCVs is the EMCV IRES, we hypothesized that DDX60’s antiviral action may be against the EMCV IRES, and not a component of HCV per se.

### DDX60 inhibits plasmid- and *in vitro* transcribed RNA-based reporters translationally driven by type II internal ribosome entry sites

The IRES of EMCV is one of four different types of commonly recognized IRESs. To determine whether DDX60 inhibits other IRES types as well, we tested the effect of DDX60 expression on two complementary reporter-based systems: a plasmid-based dual luciferase bicistronic reporter system (Honda et al., 2000; Jackson, 2013; Pelletier & Sonenberg, 1988), and a monocistronic RNA-based reporter system. The plasmid-based dual luciferase bicistronic reporter construct, once introduced into cells, is transcribed and canonically 5’-capped in the nucleus by the host cell machinery. In the resulting single bicistronic mRNA, translation of the first cistron, Renilla luciferase (Rluc), is initiated by a canonical 5’ cap mechanism, and translation of the second cistron, Firefly luciferase (Fluc), is initiated by an IRES mechanism. A stop codon separates the Rluc and Fluc genes such that Fluc can only be translated if a cap independent IRES allows for translation initiation (see **Fig. 2A, rightmost panel for schematic of transcript**). The advantage of this system is the fact that the host cell machinery provides a canonical cap at the 5’ end of the transcribed mRNA. However, since translation of Rluc and Fluc initiate from the same transcript, one must consider potential cis-acting effects when interpreting results. The RNA-based reporter system has an advantage in this regard. This system utilizes a monocistronic *in vitro* transcribed Fluc mRNA that, upon entering the cell, is immediately translated by either different IRES types or a synthetic 5’ cap analog. The use of a synthetic 5’ cap analog is a limitation in this system, although translation initiation can occur in cell free lysates similar to canonical cap structures added using a guanylyl transferase (Krieg & Melton, 1987). IRES-driven and cap-driven reporter translation are uncoupled in the RNA-based system, such that only one translation mechanism is assayed at a time. Additionally, there is no transcription or nuclear translocation, thus better simulating only the translation step of most RNA virus infections, which complete their life cycle in the cytoplasm. In summary, both reporter assays have their merits and caveats but complement each other. We chose to perform the reporter assays in HEK293T cells due to their low DDX60 expression in the absence of type I IFN stimulation (**Fig. S1A, E**) and ease of transfectability due to the absence of the DNA sensing pathway protein, STING (Sun et al., 2013). This allowed us to simulate the effects of DDX60 upregulation in the absence of endogenous DDX60 with the caveat that the exogenous DDX60 protein levels are about seven times higher than the amount of endogenous DDX60 synthesized after 48h of IFN-ß treatment (**Fig. S1E**).

**Figure 2.**
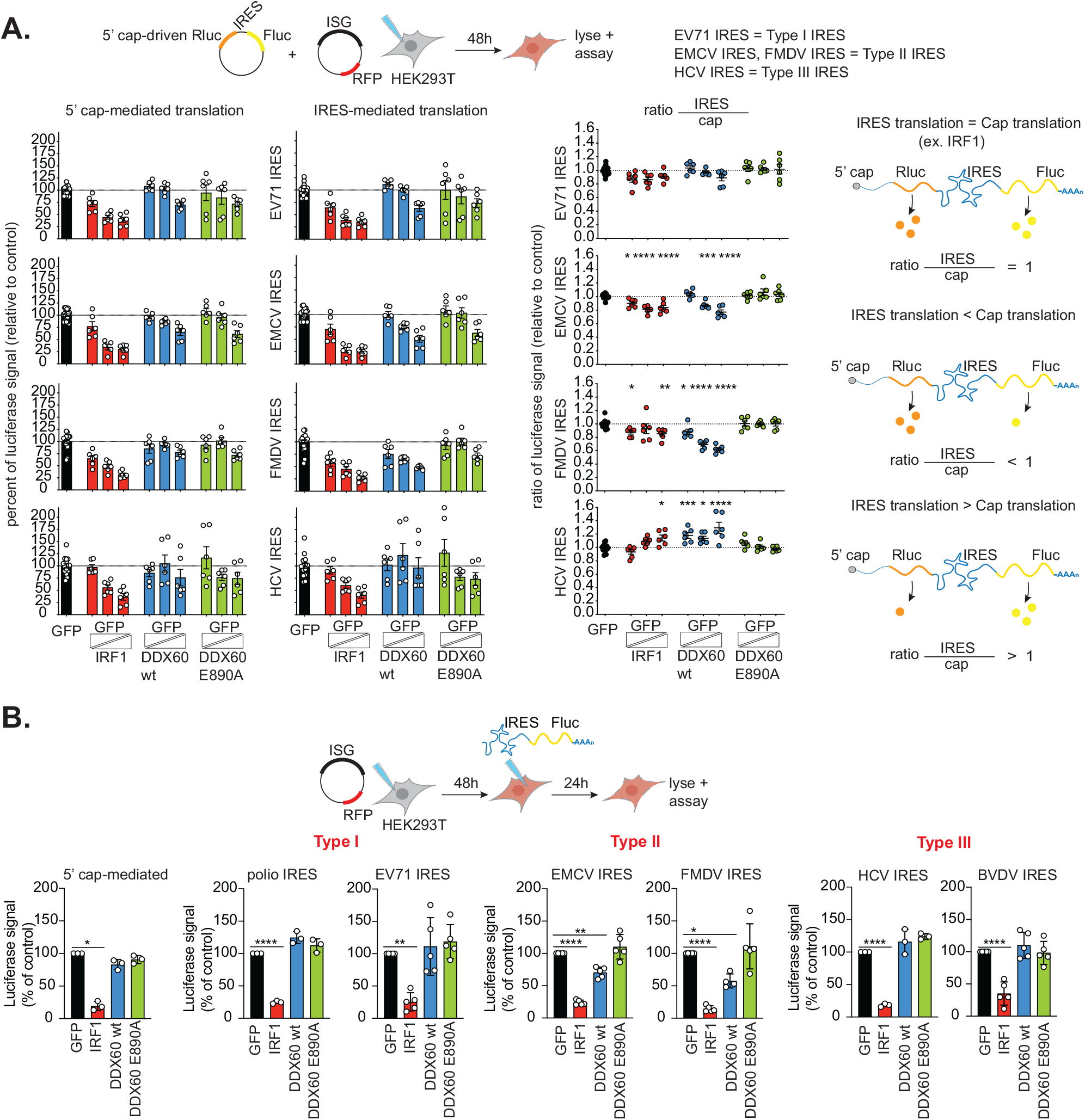
In cell reporter assay for reporters translationally driven by different internal ribosome entry sites. **(A)** Plasmid-based dual luciferase bicistronic reporter assays. HEK293T cells co-transfected with dual luciferase bicistronic reporter plasmid (Renilla luciferase (Rluc) translationally driven by a 5’ cap, and Firefly luciferase (Fluc) translationally driven by different IRESs as indicated) and GFP plasmid (negative control) or increasing amounts of IRF1 (positive control), DDX60 wt, or DDX60 E890A mutant. Total amount of DNA transfected was kept constant by supplementing transfection mixes with GFP plasmid. Luciferase units after cell lysis is plotted as a percentage of GFP transfected cells (left) and ratio of IRES Fluc units over 5’ cap Rluc units (middle). Diagram to the right of ratios plot depicts expected ratios of IRES-driven Fluc units to 5’ cap-driven Rluc units given either: equal translation of 5’ cap-driven Rluc and IRES-driven Fluc (top), greater translation of IRES-driven Fluc (middle), or greater translation of 5’ cap-driven Rluc (bottom). **(B)** RNA-based monocistronic luciferase reporter assays. HEK293T cells transfected with GFP (negative control), IRF1 (positive control), DDX60 wt, or DDX60 E890A and subsequently transfected with in vitro transcribed 5’ cap or different IRES driven Fluc mRNA constructs as indicated. Luciferase units after cell lysis is plotted as a percentage of GFP transfected cells. Raw data is shown in Figure S2B. Data shows mean ± SD for at least 3 biological replicates from experiments performed on separate days; *p < 0.05, **p < 0.01, ***p < 0.001, ****p < 0.0001 using ANOVA and Dunnett’s multiple comparison test against GFP.

In a first set of experiments, we screened for DDX60’s ability to inhibit representatives of type I, type II, or type III IRESs using the plasmid-based dual luciferase reporter system; the type IV IRES of CrPV was excluded because it has low activity in mammalian cells (Carter et al., 2008). To do so, we co-transfected HEK293T cells with dual luciferase bicistronic reporter plasmids containing different IRESs along with GFP as a negative control, increasing amounts of IRF1 as a positive control, of wild type DDX60, or a DDX60 helicase domain mutant (DDX60 E890A) while maintaining equal DNA transfection amounts by supplementing with GFP plasmid. We then analyzed Rluc and Fluc activity from cell lysates. First, we separately analyzed Rluc (cap), and Fluc (IRES) activity relative to our GFP only transfected negative control. DDX60 did not inhibit 5’ cap-driven Rluc production nor EV71 (type I) IRES- and HCV (type III) IRES-driven Fluc production but did inhibit EMCV and FMDV (type II) IRES-driven Fluc production. Additionally, our positive control, IRF1, inhibited both 5’ cap-driven Rluc production and IRES-driven Fluc production in a dose-responsive manner regardless of which IRES was present (**Fig. 2A, left**). We also observed decreases in Rluc and Fluc production for the highest levels of transfected transgenes, regardless of construct, but this may have been due to uniformly less uptake of the bicistronic reporter during transfection (**Fig. 2A, left**). We then normalized the IRES-driven Fluc activity by the 5’ cap-driven Rluc activity to account for transfection efficiencies. We hypothesized three scenarios assuming inhibitory effects at the step of translation as depicted on the right in Figure 2A: equal translation of both IRES-driven Fluc and 5’ cap-driven Rluc, decreased translation of IRES-driven Fluc compared to 5’ cap-driven Rluc, or greater translation of IRES-driven Fluc compared to 5’ cap-driven Rluc. Our positive control, IRF1, inhibited both IRES-driven Fluc production and 5’ cap-driven Rluc production equally, giving a ratio of approximately 1 (**Fig. 2A**). In contrast, DDX60 had a statistically significant dose responsive inhibitory effect on the type II IRESs of EMCV and FMDV, but not of the type I IRES of EV71 or type III IRES of HCV. This effect was lost due to DDX60 E890A mutation (**Fig. 2A**).

In the above assay, we noticed that DDX60 may have cis-acting inhibitory effects on Rluc production when Rluc is linked to EMCV IRES-driven Fluc (**Fig. 2A, left bar graphs**). Cis-acting effects may occur if mechanisms of inhibition include either RNA degradation or deterring ribosome accumulation on the entire transcript. We thus assayed *in vitro* transcribed monocistronic Fluc mRNAs that are translationally driven by either a 5’ cap analog or IRESs from type I, type II, and type III families to be able to individually assess each translation mechanism and disentangle results from potential cis-acting effects. We transfected equimolar amounts of *in vitro* transcribed Fluc mRNAs to either GFP, IRF1, DDX60 wt, or DDX60 E890A transfected cells and measured Fluc activity. Consistent with our findings from the plasmid-based system, we found that DDX60 significantly reduced translation of Fluc from mRNAs driven by the type II IRESs of EMCV and FMDV, but not by other IRES types or a 5’ cap (**Fig. 2B and Fig. S2B**). Overall, we conclude that DDX60 inhibits the type II IRES family but not the other IRES types or 5’ cap-driven translation.

Next, we asked whether other motifs or regions in DDX60 apart from the helicase motif were important for inhibiting different IRESs using our monocistronic RNA reporter system. Probing our existing panel of DDX60 mutants, we found that predicted ATP binding residues, helicase motif, and N and C-terminal extensions are important for DDX60 anti-type II IRES activity (**Fig. S2A, type II IRES panel**). We additionally found that one DDX60 mutant (domain I K791N, a presumed ATP binding/hydrolysis mutant) increased all types of IRES-driven translation but had no effect on 5’ cap-driven translation, and one DDX60 mutant (domain VI Q1321A, also a presumed ATP binding/hydrolysis mutant) uniformly decreased both 5’ cap-driven translation and all types of IRES-driven translation (except for HCV IRES) (**Fig. S2A**). The findings from this analysis of mutants correlate with the EMCV IRES-driven RFP findings and bicistronic reporter HCV findings demonstrated in figures 1B and 1C. Future biochemical and structural studies comparing these mutants with wild type DDX60 may elucidate the enzymatic activities responsible for the observed effects.

**Figure S2.**
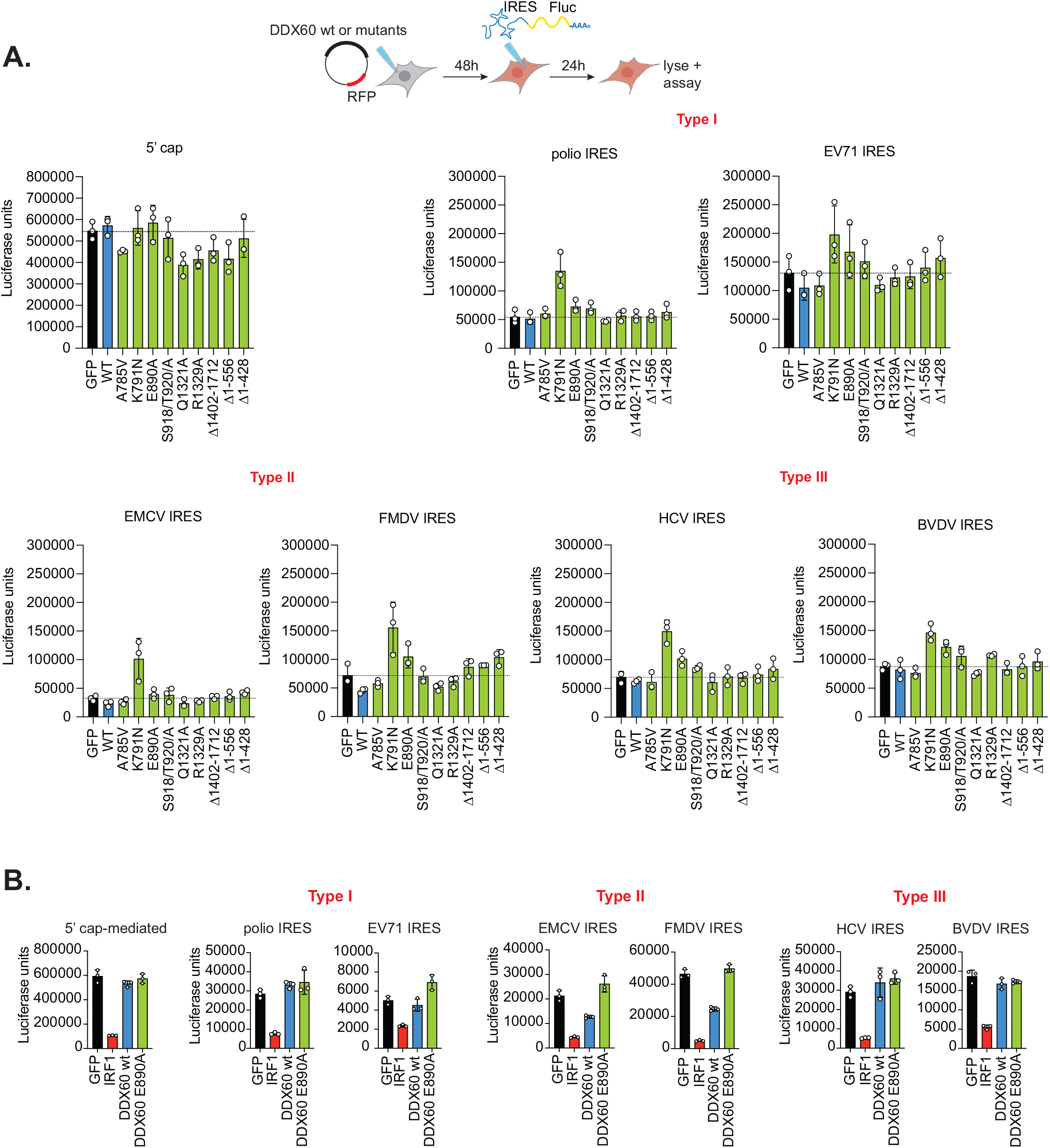
RNA reporter assays for different internal ribosome entry sites and DDX60 mutant panel. **(A)** HEK293T cells transfected with GFP (negative control), DDX60 wt, or different DDX60 mutants described in Figure 1 and subsequently transfected with in vitro transcribed 5’ cap- or different IRES-driven Fluc mRNA constructs. Mean ± SD luciferase units from lysed cells are plotted. Dotted line shows mean luciferase units from Fluc reporters transfected into GFP transfected cells. Conditions below dotted line show inhibition of Fluc production and conditions above the dotted line show enhancement of Fluc production. Data is an average of 3 biological replicates from experiments performed on separate days. **(B)** HEK293T cells transfected as in Fig. 2B. Luciferase units after cell lysis is plotted. Data shows mean ± SD for 3 technical replicates from one representative biological replicate for data shown in Fig. 2B.

### DDX60 specifically inhibits viruses that rely on type II IRES mediated translation

Thus far we demonstrated DDX60’s ability to inhibit type II IRESs using plasmid and RNA based reporters. We next sought to determine whether DDX60 can inhibit type II IRESs in the context of a virus infection. In a first set of experiments, we chose poliovirus as a representative type I IRES containing virus and EMCV a representative type II IRES containing virus, as we could work with both viruses in our BSL2 environment. We chose HeLa cells for these infection experiments, as they are highly permissive to both viruses (Jin et al., 1994; Mendelsohn et al., 1989) and express low levels of endogenous DDX60 (**Fig. S1H**). We generated HeLa cells stably expressing either wild type DDX60, DDX60 E890A, Fluc as a negative control, or IRF1 as a positive control (**Fig. S3B**). We then performed multi-cycle growth kinetics with poliovirus or EMCV to allow for DDX60’s inhibitory action to accumulate over multiple cycles of virus replication. We observed wild type DDX60 to reduce EMCV (type II IRES, **Fig. 3B**), but not poliovirus (type I IRES, **Fig. 3A**) titers relative to DDX60 E890A. The reduction in EMCV titers peaked to a two-fold reduction 24-hours post infection (hpi), and then titers increased to levels observed in our negative control and DDX60 E890A mutant expressing cells at 48 hpi, possibly owing to EMCV titers becoming high enough to overcome any inhibitory effects by DDX60 (**Fig. S3D**). Poliovirus, on the other hand, was not inhibited by DDX60 at any time point in our assay; its titers remained similar between DDX60, our negative control (Fluc), and DDX60 E890A mutant expressing cells at all time points even while our positive control, IRF1 reduced poliovirus titers starting at 24 hpi (**Fig. S3C**). Broad transcriptional regulators like IRF1 activate a cascade of other antiviral genes to potently inhibit titers of multiple viruses by several orders of magnitude (Schoggins et al., 2011). Unlike IRF1, we anticipated DDX60 to have specific one-on-one antiviral activity, possibly explaining its more modest reduction of EMCV titers compared to IRF1.

**Figure 3.**
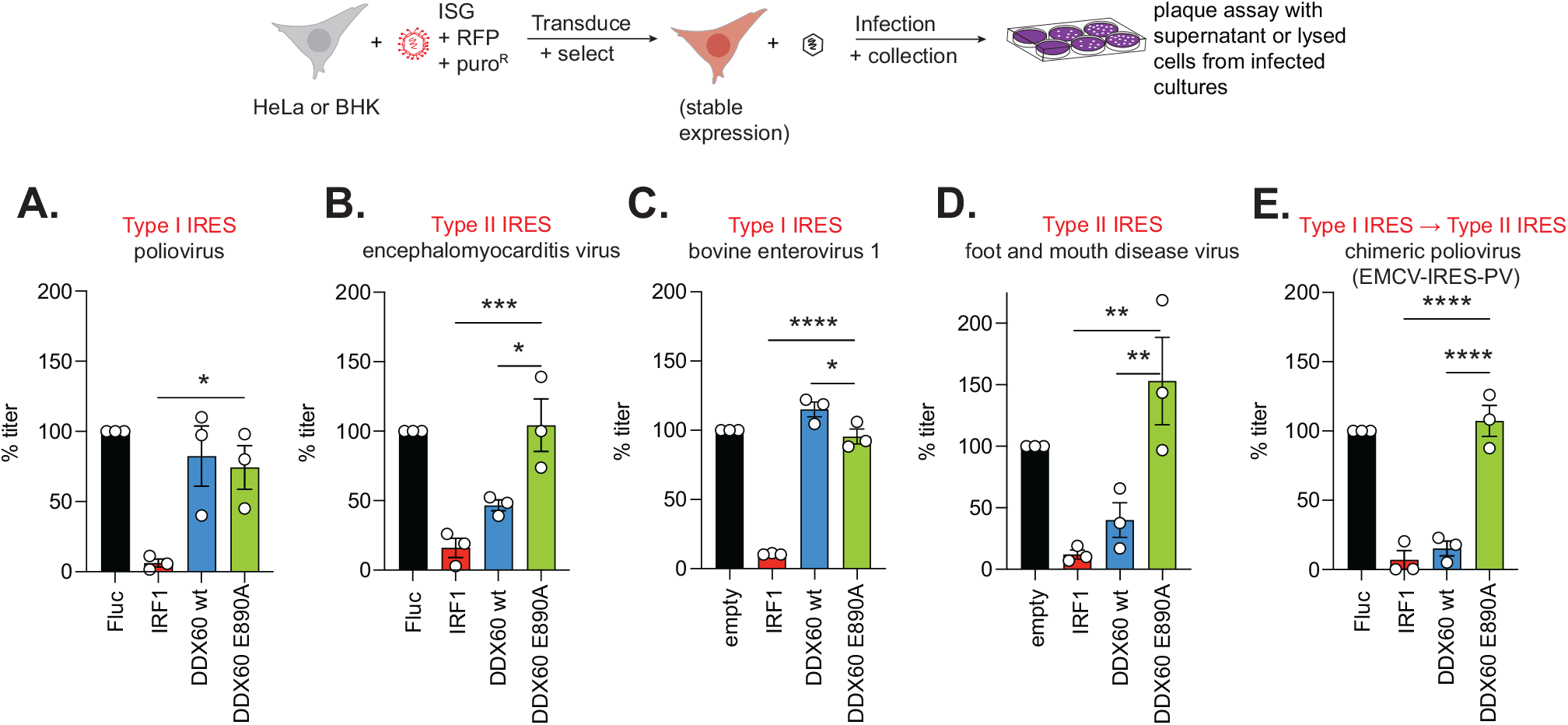
Viral replication assays for a panel of IRES containing viruses. **(A, B, E)** Multi-cycle infection assays with type I and type II IRES containing viruses. HeLa cells stably expressing Firefly luciferase (Fluc) (negative control), IRF1 (positive control), DDX60 wt, or DDX60 E890A and infected with **(A)** poliovirus, **(B)** encephalomyocarditis virus, or (**E**) a chimeric poliovirus with the poliovirus IRES replaced with the IRES of EMCV (EMCV-IRES-PV) at MOI 0.001. Data shows mean ± SEM percent infectious titers relative to Fluc from supernatants collected 24 hours post infection (hpi) (EMCV) or 48 hpi (poliovirus and EMCV-IRES-PV). Titers determined via plaque assay on HeLa cells. **(C, D)** Single-cycle infection assays with type I and type II IRES containing viruses. BHK-J cells stably expressing empty vector (negative control), IRF1 (positive control), DDX60 wt, or DDX60 E890A and infected with **(C)** bovine enterovirus-1, or **(D)** foot and mouth disease virus at MOI 1. Data shows mean ± SEM of infectious titers from lysed cells relative to Fluc. PFU data is shown in Figure S3. Titers determined via plaque assay on BHK-21 clone 13 cells. Data is from at least 3 biological replicates from experiments performed on separate days; *p < 0.05, **p < 0.01, ***p <0.001, ****p<0.0001 using ANOVA and Dunnett’s multiple comparison test against DDX60 E890A.

Next, we performed infection experiments with another type II IRES-containing virus, FMDV. In the US, these experiments are only possible at Plum Island Animal Disease Center’s enhanced BSL3 facility. We compared DDX60 action against FMDV to that of action against bovine enterovirus 1 (BEV-1), which carries a type I IRES. We performed these infections in baby hamster kidney (BHK) cells, as human cells are not permissive to FMDV or BEV-1 (Mowat & Chapman, 1962; Ruiz-Sáenz et al., 2009). BHK-J cells stably expressing either a negative control empty vector, positive control IRF1, wild type DDX60, or DDX60 E890A were infected with either BEV-1 or FDMV at an MOI of 1 and progeny virus harvested at 5 hpi. This allowed us to observe the magnitude of IRES inhibition by DDX60 on infectious titers after a single round of virus replication. Consistent with our findings using EMCV and poliovirus, we detected a statistically significant reduction in titers of FMDV (type II IRES, **Fig. 3D**) but not BEV-1 (type I IRES, **Fig. 3C**). These results demonstrated that DDX60 inhibits viruses that rely on type II IRES-driven translation.

Next, we tested whether a type II IRES is sufficient to confer sensitivity to DDX60. We thus generated a chimeric poliovirus replacing its endogenous type I IRES with the type II IRES of EMCV (EMCV-IRES-PV). We first characterized EMCV-IRES-PV replication in comparison to poliovirus and EMCV. First, we noticed that EMCV-IRES-PV generated smaller plaques compared to poliovirus as observed for a similar chimeric virus generated previously (Alexander et al., 1994), data not shown). In multi-cycle replication kinetics, EMCV-IRES-PV started producing detectable infectious particles in HeLa cells between 8 and 24 hpi. Its titers peaked to approximately 10^9^ PFU/mL at the end point of our assay, 48 hpi (**Fig. S3A**). This replication dynamic resembled that of poliovirus rather than EMCV. EMCV produced infectious particles of 10^6^ PFU/mL after just 8 hpi with peak titers of approximately 10^10^ PFU/mL at 24 hpi (**Fig. S3A**). EMCV-IRES-PV and poliovirus both produced titers 10- to 100-fold lower than EMCV for most of the experiment, but eventually reached similar titers as EMCV 48 hpi (**Fig. S3A**). Next, we analyzed DDX60’s ability to inhibit EMCV-IRES-PV. In contrast to its parental poliovirus strain, which was resistant to DDX60 (**Fig. 3A**), DDX60 reduced EMCV-IRES-PV titers beginning at 24 hpi, reaching a ten-fold titer reduction by the end of our assay at 48 hpi (**Fig. 3E, S3E**). Remarkably, this inhibitory effect is similar in magnitude to our positive control, IRF1, and is lost due to DDX60 E890A mutation. Together, our data demonstrate that reliance on type II IRES-driven translation is sufficient to allow DDX60-mediated inhibition of virus infection.

**Figure S3.**
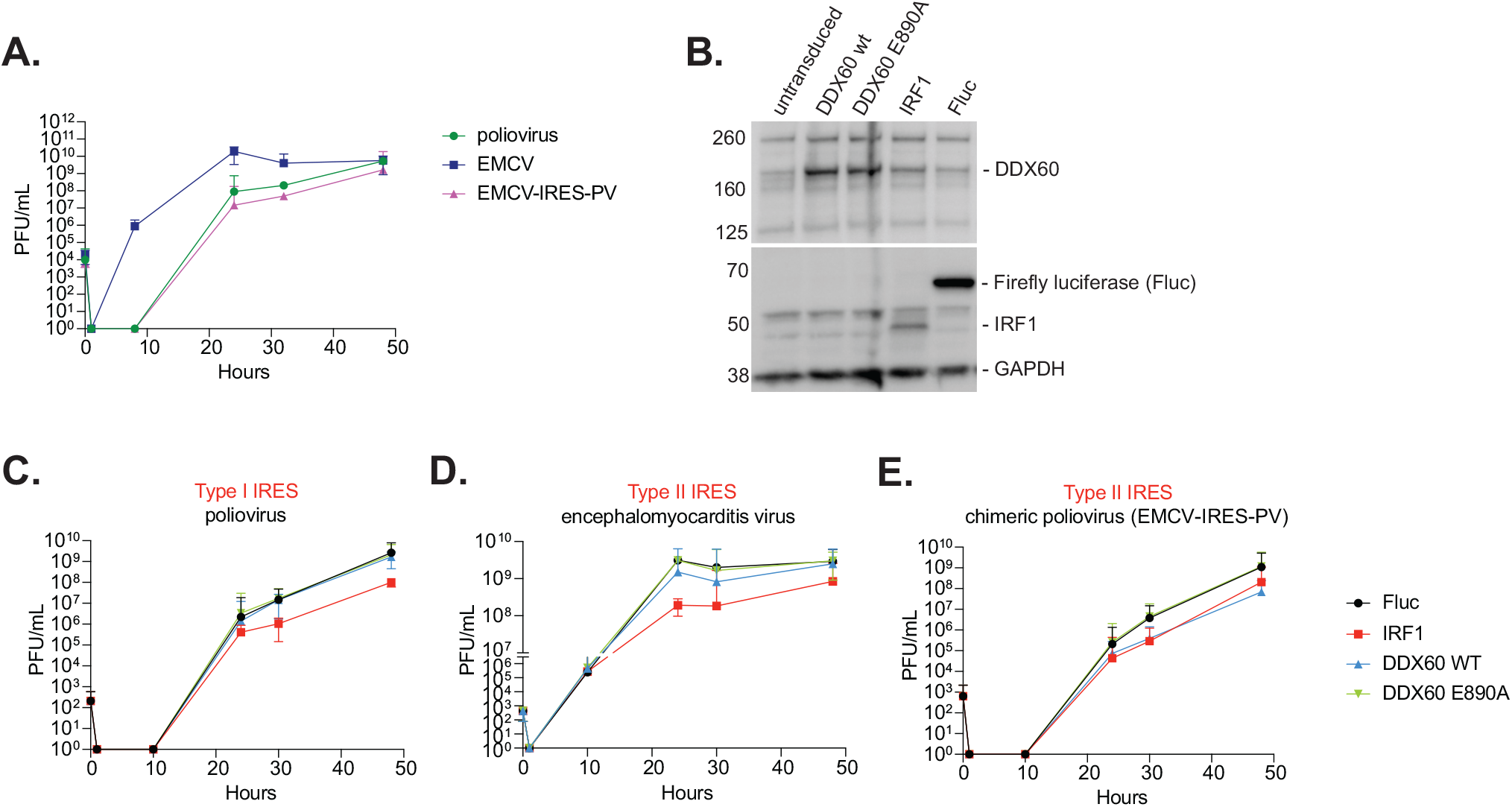
Growth kinetics of different internal ribosome entry site containing viruses in the presence of DDX60. **(A)** Growth kinetics of poliovirus, encephalomyocarditis virus (EMCV), or a chimeric poliovirus with the poliovirus IRES replaced with the IRES of EMCV. HeLa cells were infected with the three viruses at MOI 0.001 and supernatants from infected cultures were collected 0, 1, 8, 24, 32 or 48 hpi. Virus in supernatants was then titered on HeLa cells. Error bars represent mean ± 95% confidence interval. Data is representative of at least 2 biological replicates from experiments performed on separate days. **(B)** Assessment of exogenous DDX60 expression in HeLa cells. HeLa cells were transduced with lentivirus carrying either DDX60 wt, DDX60 E890A, IRF1, or Firefly luciferase (Fluc) together with a puromycin resistance gene and selected with puromycin. Cell lysates were analyzed using western blot for DDX60, Fluc, IRF1, and GAPDH (loading control) protein products. **(C, D, E)** Time course of antiviral assays with panel of IRES containing viruses. HeLa cells stably expressing Fluc (negative control), IRF1 (positive control), DDX60 wt, or DDX60 E890A were infected with **(C)** poliovirus, **(D)** EMCV, or **(E)** a chimeric poliovirus at a MOI of 0.001. Supernatants from infected cultures were collected at either 0, 1, 8, 24, 32, or 48 hpi and titered on HeLa cells via plaque assay. Error bars represent mean ± 95% confidence interval. Data is representative of at least 2 biological replicates from experiments performed on separate days.

### Abundance of type II IRES containing mRNAs is unchanged in the presence of wild type or mutant DDX60

Next, we sought to decipher the mechanism of type II IRES inhibition by DDX60. Our first approach was to purify DDX60 to analyze its structure, ATP binding and hydrolysis activities, and RNA binding and unwinding activities. We expressed full-length ∼200 kDa DDX60 with various N-terminal tags in both yeast (Pichia X-33) and insect cells (Sf9). Expression levels were similar in both systems, and Sf9 cells were chosen for ease of protein extraction. Use of a twin-step-tag and strep-tactin XT resin allowed for an efficient first one-step protein purification, followed by a second purification through a heparin column. However, the final size exclusion chromatography step was hampered by protein aggregation, which resulted in low yields. Attempts to solubilize the protein with differing salt and glycerol concentrations were unsuccessful, thus hampering any further biochemical studies. Additionally, our findings that both N- and C-terminal extensions of DDX60 were necessary for its antiviral and anti-IRES activity (Fig. 1C, Fig. S2A) precluded us from using DDX60 helicase domains alone. We were thus limited to our cell-based assays for the remainder of this study.

We first hypothesized that DDX60 inhibits type II IRES translation by decreasing the abundance of mRNAs with a type II IRES. DDX60 was previously shown to be most closely related to a family of helicases called superkiller-2 (Ski-2)-like helicases (Goubau et al., 2015; Miyashita et al., 2011). Ski-2 is an RNA helicase originally discovered in yeast that is important for degrading satellite dsRNA from L-A double-stranded RNA virus and for general 3’ to 5’ degradation of yeast mRNAs (Anderson & Parker, 1998; Widner & Wickner, 1993). To test our hypothesis, we utilized our Fluc mRNAs translationally driven by a 5’ cap analog or different IRESs. Using this system had the following advantages over virus infection experiments: first, the reporter mRNAs are meant to only replicate the translation step of the virus, allowing us to omit other steps of the viral life cycle when interpreting results; second, IRES containing viruses like poliovirus, EMCV, and FMDV are known to dampen host mRNA translation to favor viral mRNA translation (Lee et al., 2017; Stern-Ginossar et al., 2019; Yamamoto et al., 2017), introducing many confounding variables; third, all the reporter mRNAs have the same sequence apart from the IRESs, taking away the need to consider coding sequence differences in the mRNAs when interpreting results; and lastly, side-by-side luciferase assays can corroborate any findings from measuring relative RNA abundance using RT-qPCR. A caveat of this system, however, is that transfected mRNA can both be translated in the cytoplasm or remain untranslated in endosomes, where it is either degraded or trafficked back to the cytoplasm (Johannes & Lucchino, 2018). While we could not prevent the latter possibility, we reasoned that the reporter mRNAs would all be similarly distributed in the cytoplasm versus the endosome as they are distinguished only by the presence of a 5’ cap and the IRESs they carry. This would make each mRNA type present in the cytoplasm equally susceptible to possible DDX60-mediated degradation. We thus transfected HEK293T cells to express either wild type DDX60 or DDX60 E890A, subsequently transfected with equimolar amounts of Fluc reporter RNAs translationally driven by either a 5’ cap analog, type I IRESs, type II IRESs, or type III IRESs, and analyzed cell lysates for luciferase signal and reporter mRNA content in parallel. We found that while wild type DDX60 reduced Fluc translation from type II IRES-driven mRNAs compared to DDX60 E890A, the relative RNA abundance of type II IRES-driven mRNAs between wild type DDX60 and DDX60 E890A in the same cells were equivalent (**Fig. 4**). For the other IRES types and 5’ cap-driven translation, wild type DDX60 and DDX60 E890A transfected cells had both equal levels of Fluc translation and mRNA abundance (**Fig. 4**). This indicated to us that while DDX60 can reduce protein synthesis from type II IRES containing mRNAs, it is does not reduce the abundance of such mRNAs, suggesting some other mechanism of inhibition other than decreasing mRNA abundance.

**Figure 4.**
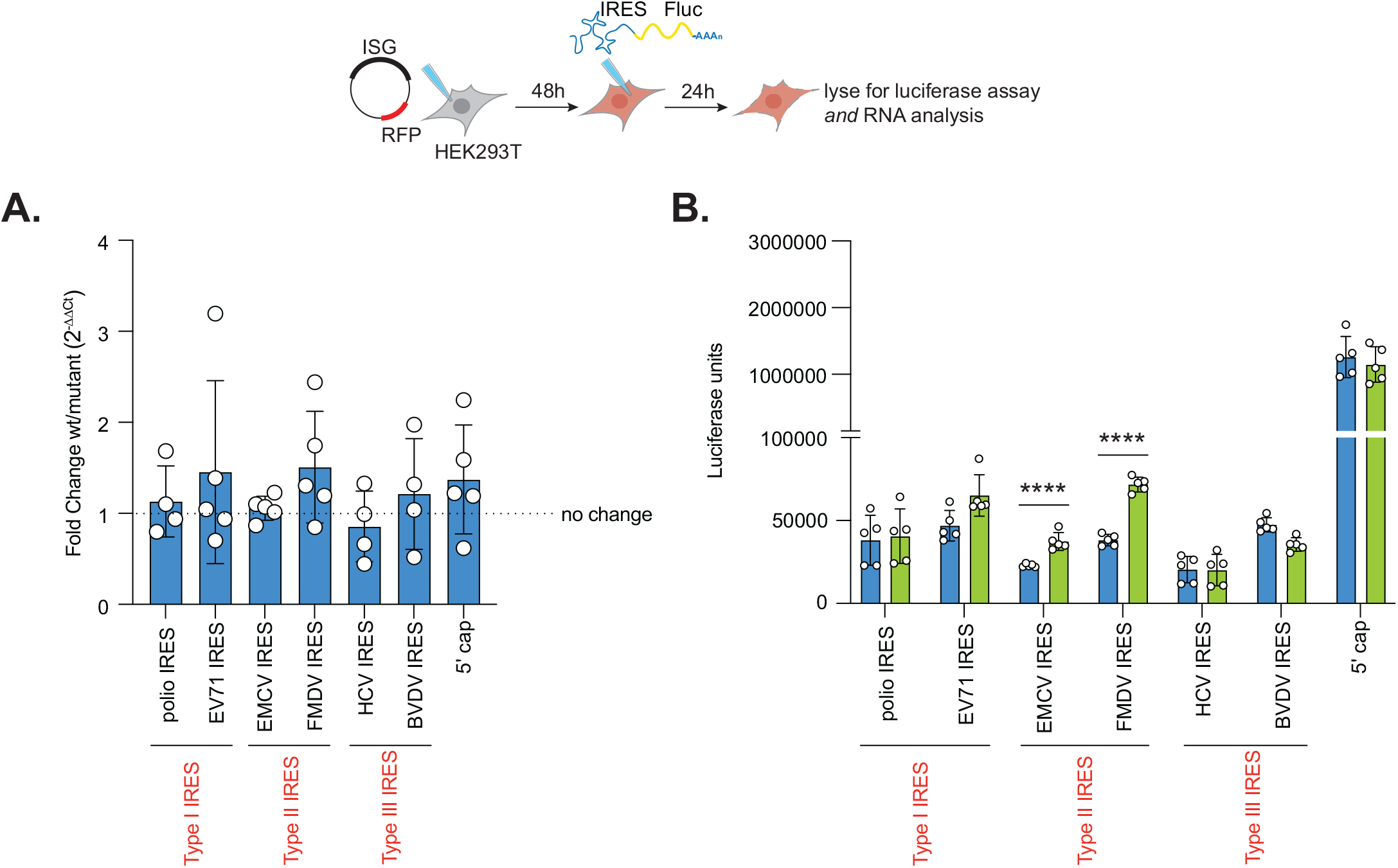
Assessing mRNA reporter abundance for reporters translationally driven by different internal ribosome entry sites. HEK293T cells transfected with DDX60 wt or DDX60 E890A (negative control) and transfected with in vitro transcribed 5’ cap- or IRES-driven Fluc mRNA constructs as indicated and lysed 24 hours later. (**A**) Abundance of luciferase reporter mRNAs assayed using RT-qPCR. **(B)** IRES- or cap-driven translation from the same samples assayed in parallel using luciferase assay. Mean ± SD luciferase units are plotted for at least 3 biological replicates from experiments performed on separate days; ****FDR < 0.01% (p<0.0001) using unpaired t-test with Welch correction and Benjamini and Yekutieli correction for multiple testing comparing DDX60 wt versus DDX60 E890A transfected cells.

### DDX60 does not enhance IFN signaling to inhibit type II IRESs

Another potential mechanism of DDX60 antiviral action is indirect, through induction of IFN. Previous publications proposed that DDX60 physically interacts with RIG-I and synergistically enhances IFN production and thus downstream IFN signaling upon recognition of a viral pathogen associated molecular pattern (PAMP) (Miyashita et al., 2011; Oshiumi et al., 2015). However, a second publication could not observe any physical interaction between DDX60 and RIG-I or enhancement of IFN signaling (Goubau et al., 2015). We thus tested whether the DDX60- and RIG-I-mediated enhancement of downstream IFN signaling contributed to its anti-type II IRES activity. To do so, we used a HEK293 reporter cell line that encodes an interferon sensitive response element (ISRE)-driven Firefly luciferase (ISRE:Fluc) in its genome. Fluc activity was minimal when cells were treated with PBS but increased 10-fold upon poly(I:C) transfection and approximately 400-fold upon 500 U/mL IFN-ß treatment (**Fig. S4A**). Next, we transfected RIG-I into our HEK293 reporter cell line and either treated with PBS or transfected with poly(I:C) to trigger signaling through RIG-I and induce IFN expression. RIG-I expression alone caused ISRE:Fluc activity to increase approximately 150-fold compared to untransfected, PBS treated controls (**Fig. S4A**). Upon poly(I:C) transfection in addition to RIG-I expression, ISRE:Fluc activity further increased to approximately 300-fold compared to untransfected, PBS treated controls, albeit this further increase was not statistically significant (**Fig. S4A**). We then proceeded to transfect increasing amounts of wild type DDX60 or DDX60 E890A in the presence of increasing amounts of transfected RIG-I. To keep the total amount of DNA transfected per condition equal, we supplemented transfection mixes with GFP plasmid. We verified that both DDX60 and RIG-I were expressed using a western blot (**Fig. S4B**). Both wild type DDX60 and DDX60 E890A transfection in combination with RIG-I, but without the presence of poly(I:C), slightly increased ISRE:Fluc activity from 150-fold to approximately 200-fold compared to untransfected, PBS treated controls, albeit not statistically significant (**Fig. S4A**). Adding transfected poly(I:C) to both wild type DDX60 or DDX60 E890A and RIG-I transfected cells did not further increase ISRE:Fluc activity any more than RIG-I and poly (I:C) only transfected cells (**Fig. S4A**). This trend remained even when increasing wild type DDX60, DDX60 E890A, or RIG-I transfection levels. If enhanced downstream IFN signaling explained DDX60 anti-type II IRES activity, we would expect any enhancement in ISRE:Fluc activity to be diminished due to the DDX60 E890A mutation. Given that we did not observe such a phenomenon suggested that downstream IFN signaling enhancement does not account for DDX60 anti-type II IRES activity.

**Figure S4.**
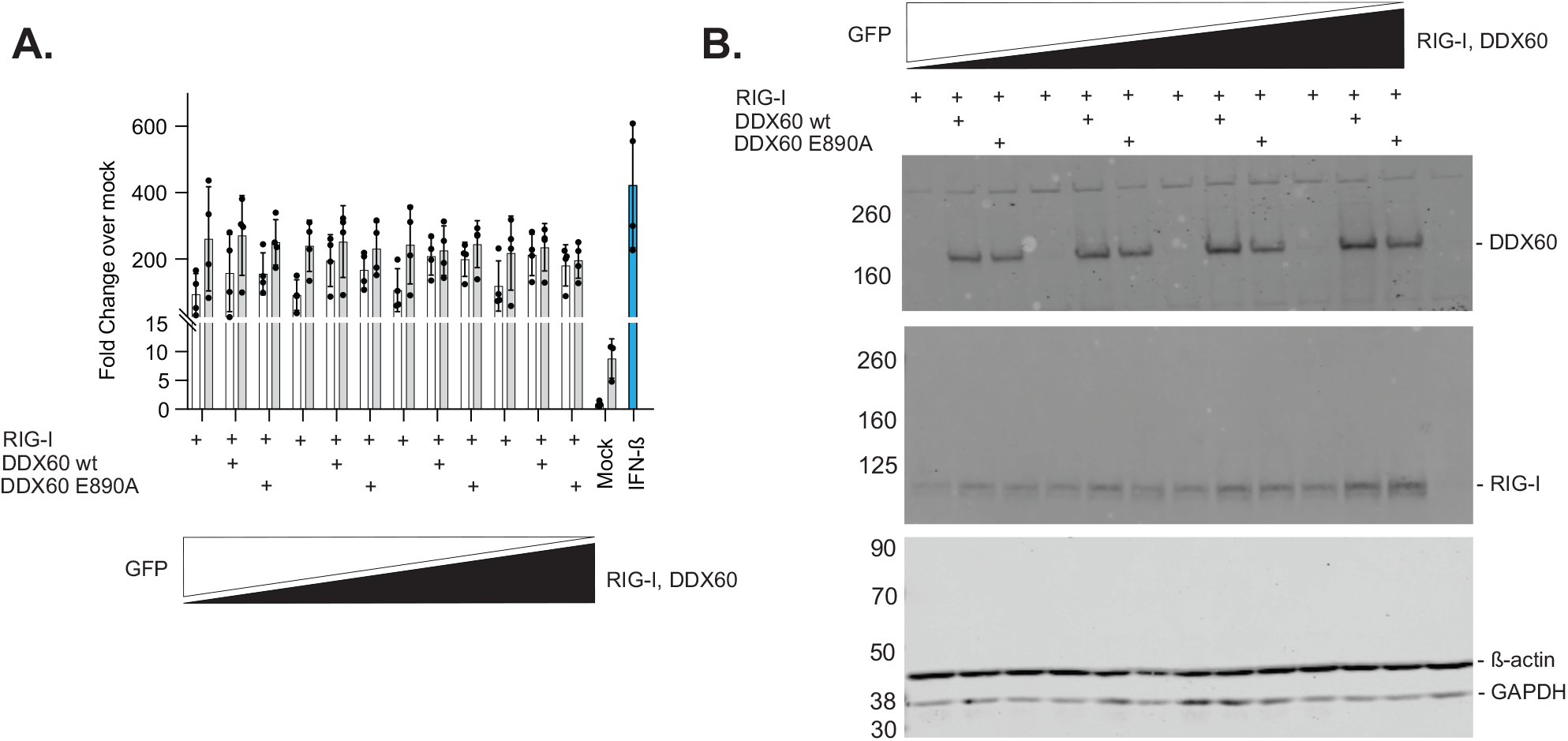
ISRE reporter assays in the presence of DDX60. HEK293T cells stably expressing a type I IFN sensitive response element (ISRE) driven Fluc gene were transfected with increasing amounts of RIG-I, RIG-I in combination with DDX60 wt, or RIG-I in combination with DDX60 E890A while supplementing with GFP plasmid to equalize the total amount of DNA transfected. Cells were then treated with either PBS (negative control), transfected with LMW poly(I:C), or treated with IFN-ß (positive control). Cells were subsequently used for a luciferase assay to assess ISRE activity **(A)** or western blot for analysis of DDX60, RIG-I, ß-actin (loading control), or GAPDH (loading control) protein products **(B)**. Mean ± SD relative luciferase units (RLU) are plotted in (A). Data is representative of at least 3 biological replicates from experiments performed on separate days.

### DDX60 binds RNA nonspecifically

Another possible mechanism of selective DDX60 antiviral action is through specific binding to the type II IRES and enacting a steric hinderance effect. Previous studies demonstrated that the closely related helicase, Ski-2, and the core helicase domains of DDX60 bind both single-stranded and double-stranded RNA with equal affinities, suggesting that these helicases can bind diverse RNA substrates (Halbach et al., 2012; Miyashita et al., 2011). Likewise, structural studies of several DEAD-box RNA helicases suggest that their interaction with RNA is structure-rather than sequence-dependent due to their interaction with the RNA sugar-phosphate backbone (Schütz et al., 2010; Sengoku et al., 2006). We tested if DDX60 binds type II IRES RNA preferentially as opposed to type I IRES RNA, type III IRES RNA, or 5’ capped RNA. We devised two pulldown strategies: one pulling down RNA and detecting bound DDX60, and the other pulling down DDX60 and detecting bound RNA.

First, we generated biotin-UTP labeled RNA probes that can be precipitated using streptavidin coated beads and subsequently assayed for bound proteins, such as DDX60, using western blot. A previous study used this system to identify far upstream element-binding protein 1 (FBP1) bindings sites in the EV71 IRES (Hung et al., 2016). Our panel of probes included IRESs of poliovirus (type I), EMCV (type II), HCV (type III), and 5’ capped Fluc RNA, as well as matching unlabeled probes as negative controls. To test whether DDX60 associates with these RNA sequences, we incubated cell lysates from DDX60 expressing HEK293T cells with biotin-UTP labeled or unlabeled probes. We first tested for a protein known to specifically interact with type I and type II IRESs, but not type III IRESs, the ITAF Polypyrimidine tract-binding protein 1 (PTPB1). We found PTPB1 to be enriched upon precipitation of biotin-UTP labeled poliovirus IRES and EMCV IRES probes (type I and type II, respectively) compared to matched unlabeled probes but saw no enrichment between labeled and unlabeled probes for HCV IRES and 5’ capped Fluc RNA (**Fig. S5A, left blot**). Some nonspecific PTPB1 binding observed is attributed to PTPB1 binding to streptavidin beads alone (**Fig. S5A, left blot**). Additionally, we recapitulated FBP1 binding to EV71 IRES as previously reported (Hung et al., 2016) (**Fig. S5A, middle blot**). However, when analyzing DDX60, we found DDX60 to be equally present when precipitating biotin-UTP labeled or unlabeled probes for all IRES types and 5’ capped Fluc (**Fig. S5A, right blot**). Unlike PTPB1, this nonspecific precipitation of DDX60 is not due to DDX60 binding to streptavidin beads as DDX60 does not show any binding to streptavidin beads alone (**Fig. S5A, left blot**). This led us to favor the conclusion that DDX60 is a “sticky,” non-specific RNA binder with the caveat that *in vitro* RNA binding does not necessitate binding in cells.

We next validated these findings with the converse strategy. We expressed wild type DDX60 in HEK293T cells and subsequently transfected either 5’ cap-driven Fluc mRNA or EMCV IRES-driven Fluc mRNA. As a negative control, we transfected 5’ cap-driven Fluc mRNA or EMCV IRES-driven Fluc mRNA without expressing DDX60. We then immunoprecipitated using either a DDX60 targeting antibody or isotype control IgG and analyzed Fluc mRNA quantities by RT-qPCR. We found both 5’ cap-driven Fluc mRNA and EMCV IRES-driven Fluc mRNA to be enriched upon DDX60 immunoprecipitation in our DDX60 expressing samples (**Fig. S5B, right**). In contrast, there was no enrichment of these mRNAs in samples with undetectable DDX60 expression (**Fig. S5B, right**). This provided additional evidence that DDX60 non-specifically binds both 5’ capped and EMCV IRES containing mRNAs *in vitro*. Overall, our RNA bindings assays led us to conclude that DDX60 does not distinguish type II IRES containing mRNAs by differential RNA binding.

**Figure S5.**
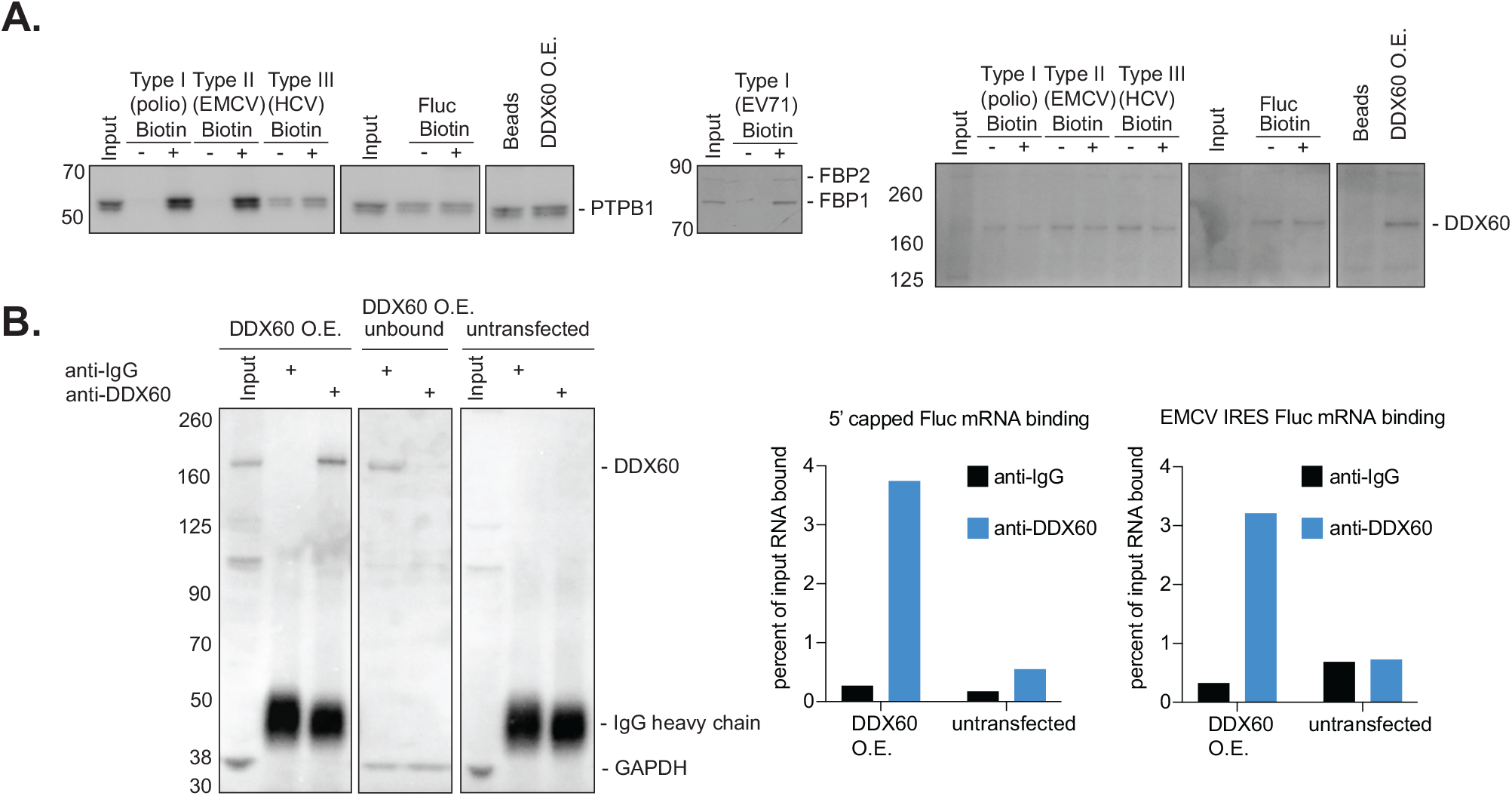
Probing for a physical interaction between DDX60 and different IRES sequences. **(A)** RNA-protein binding assays using biotin labeled RNA probes and streptavidin coated beads. DDX60 wt expressing HEK293T cell lysates were combined with either biotin labeled or unlabeled (negative control) IRES RNA sequences (IRESs from poliovirus, EV71, EMCV, or HCV) or 5’ capped Fluc RNA sequence (negative control). After allowing for protein and RNA binding, streptavidin coated beads were used to pull down RNA. After washing, RNA bound proteins were visualized using western blot. Input or DDX60 overexpression (O.E) lanes show DDX60 and other interrogated RNA binding proteins before incubation of cell lysates with interrogated RNAs. Comparisons for enrichment of RNA binding is made by comparing unlabeled and biotin labeled lanes. Visually equivalent band intensities between unlabeled and biotin labeled lanes signify non-specific binding between an RNA and interrogated protein. Lane labeled beads represents amount of protein that binds to streptavidin coated beads in the absence of any interrogated RNA. RNA binding proteins interrogated include Polypyrimidine Tract Binding Protein 1 (PTPB1) (positive control for binding type I and type II IRESs specifically), Far upstream element binding protein 1 (FBP1) (binds EV71 IRES), and DDX60. Data is representative of at least 2 biological replicates from experiments performed on separate days. **(B)** Protein-RNA binding assays using immunoprecipitated DDX60 and RT-qPCR for Fluc mRNAs. HEK293T cells transfected with DDX60 wt or left untransfected were transfected with either 5’ cap- or EMCV IRES-driven Fluc mRNA. Cell lysates were then subjected to immunoprecipitation using either DDX60 targeting antibody or isotype control IgG. Bound mRNA was detected using RT-qPCR against Fluc. Efficiency of immunoprecipitation is visualized using western blot against DDX60. Bound mRNA is quantified as a percent of input RNA post-immunoprecipitation with either IgG or anti-DDX60 antibody in either DDX60 expressing or untransfected cells. Data is representative of 2 separate immunoprecipitations.

### DDX60 impedes polysome occupancy on type II IRES containing mRNAs

Our RNA abundance assay suggested that while DDX60 does not decrease the abundance of type II IRES containing mRNAs, it still diminishes translation from the IRES as seen by the decrease in Fluc protein synthesis (**Fig. 4**). Such diminished translation may be a result of DDX60 reducing ribosome binding to the mRNAs. To test changes in ribosome binding we performed polysome profiling. Polysomes are found on heavily translated mRNAs and are a result of multiple ribosomes translating the same mRNA (Viero et al., 2015; Warner et al., 1962; Wettstein et al., 1963). Polysome associated mRNA can be isolated by ultracentrifugation through a sucrose gradient and subsequent fractionation. The more heavily translated an mRNA is, the higher the abundance of that mRNA in higher fraction numbers (Panda et al., 2017).

We first asked whether DDX60 has a global effect on translation by comparing the global polysome profile in untransfected or wild type DDX60 expressing HEK293T cells. We found that the overall distribution of mRNA in the different polysome fractions were similar between DDX60-expressing and control untransfected cells (**Fig. S6A**), suggesting that DDX60 does not affect cellular translation globally.

We next hypothesized that expression of wild type DDX60 would shift the abundance of type II IRES containing mRNAs to lower fraction numbers but leave the abundance of a non-type II IRES containing mRNAs more concentrated in higher fraction numbers. To test this hypothesis, we expressed wild type DDX60 or our negative control, DDX60 E890A, in HEK293T cells. We transfected these cells with equimolar amounts of either 5’ cap-driven or EMCV IRES-driven Fluc mRNA. Additionally, we treated duplicate samples of DDX60 E890A plus 5’ cap-driven Fluc mRNA transfected cells and DDX60 E890A plus EMCV IRES-driven Fluc mRNA transfected cells with puromycin as positive controls for translation inhibition by disrupting polysomes (**Fig. 5A**) (Kudla & Karginov, 2016). As expected, puromycin treated samples showed mRNA accumulation in lower fraction numbers (**Fig. S6B**).

**Figure 5.**
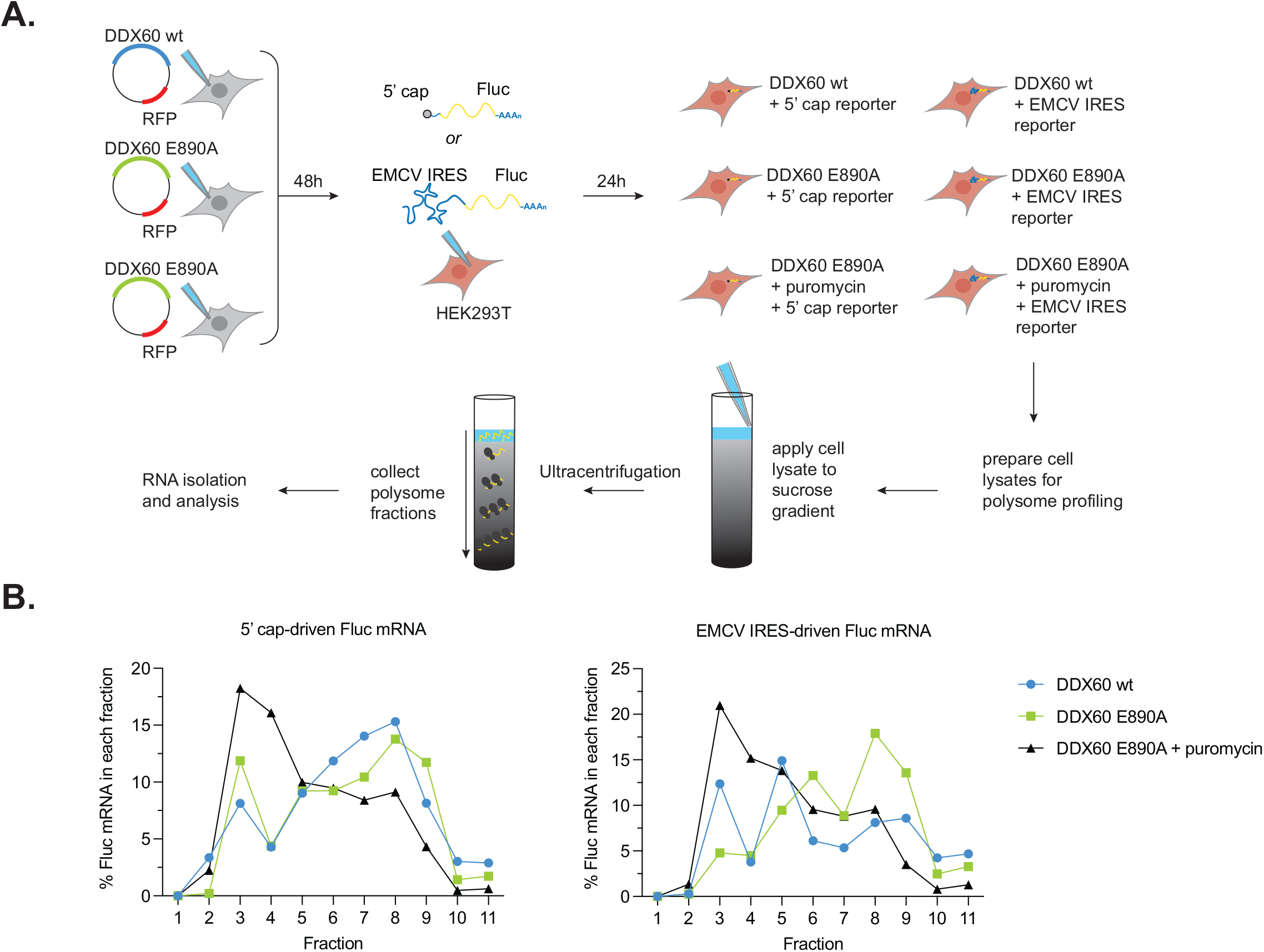
Polysome occupancy in the presence of DDX60. **(A)** Schematic of polysome profiling strategy. HEK293T cells were transfected with DDX60 wt or DDX60 E890A (negative control). 48 hours post transfection, cells were transfected with in vitro transcribed 5’ cap- or EMCV IRES-driven Fluc mRNA constructs for 24 hours. Duplicate samples were treated with 200 μM puromycin for 20 minutes as positive controls for decrease in polysomes. Cells were treated with 100 μg/mL of cycloheximide for 15 minutes to arrest polysomes and subjected to polysome profiling by ultracentrifugation through 15%-50% sucrose gradients. Amount of Fluc reporter mRNA from polysome fractions was determined by RT-qPCR. **(B)** Effect of DDX60 on polysome occupancy of 5’ cap (left) or EMCV IRES (right) driven Fluc mRNA. Shown is the mean percent of Fluc mRNA in each of the fractions from 2 biological replicates from experiments conducted on separate days. Full polysome profiles are shown in Figure S6.

We then proceeded to analyze the distribution of Fluc reporter mRNA in the different polysome fractions for each condition by RT-qPCR. Puromycin treatment shifted both 5’ cap- and EMCV IRES-driven Fluc mRNAs to lower fraction numbers, demonstrating decreased ribosome accumulation on the mRNA reporters. Further we found that wild type DDX60 expression had no effect on 5’ cap-driven Fluc mRNA distribution within fractions as it closely followed the distribution seen in DDX60 E890A expressing cells (**Fig. 5B, left**). DDX60 E890A expressing cells showed no change in the distribution of EMCV IRES-driven Fluc mRNA compared to 5’ cap-driven Fluc mRNA distribution but adding puromycin shifted the EMCV IRES-driven Fluc mRNA toward lower fraction numbers (**Fig. 5B**). Finally, DDX60 wild type expression shifted EMCV IRES-driven Fluc mRNA toward lower fraction numbers (**Fig. 5B, right**), demonstrating that DDX60 selectively decreases polysome occupancy on type II IRES-driven mRNAs.

Altogether our results show that the ISG, DDX60 reduces protein synthesis driven by type II IRESs via decreasing polysome binding to type II IRES-driven mRNAs (**Fig. 6**).

**Figure 6.**
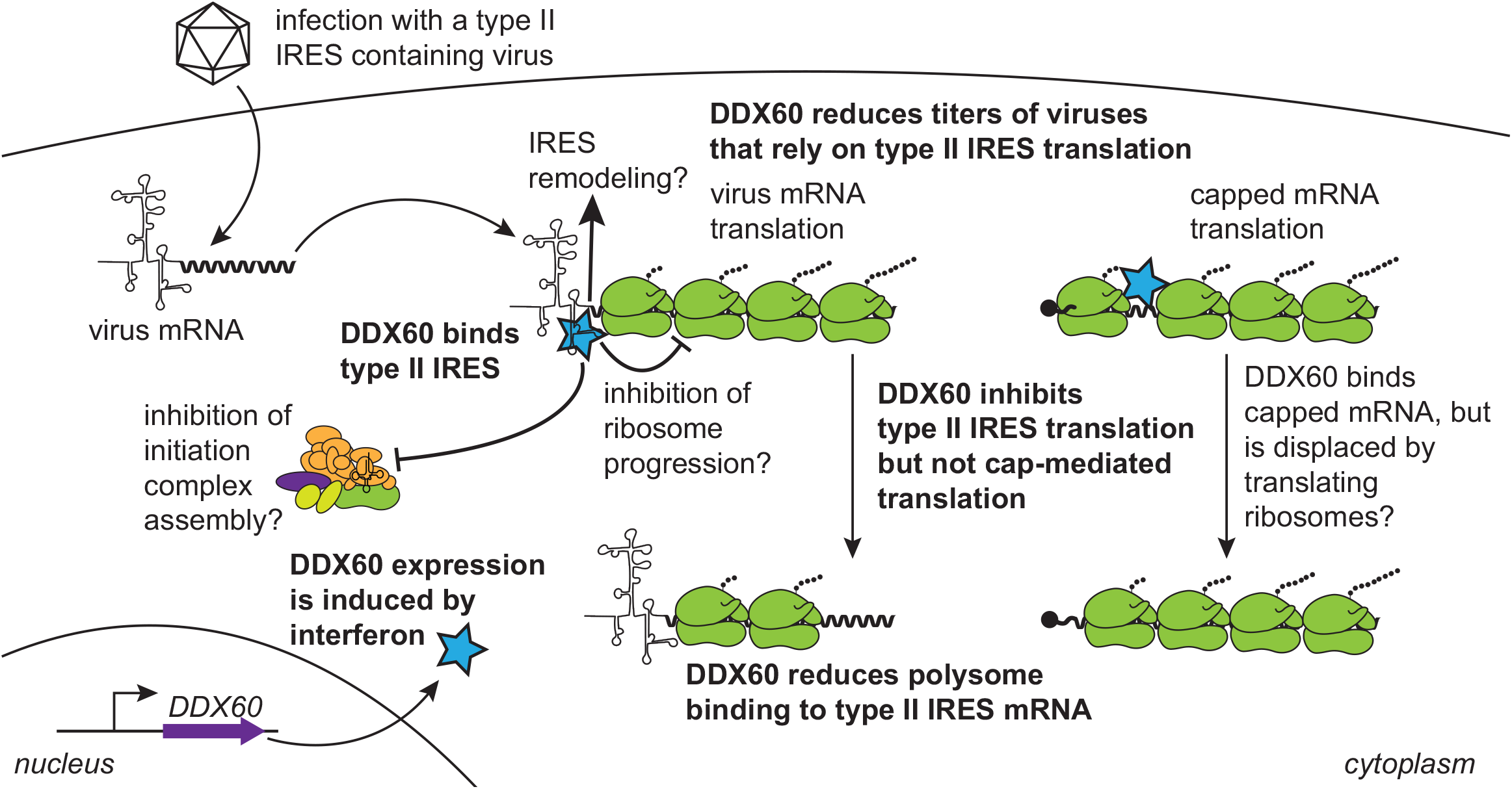
Proposed mechanisms for inhibition of type II IRES mRNA translation by DDX60. Upon entry of a type II IRES containing RNA virus, the viral mRNA can be directly translated by the host protein synthesis machinery. Virus infection also promotes the expression of interferon (IFN) and other pro-inflammatory cytokines leading to the production of an array of interferon-stimulated genes (ISGs), among which is DDX60. DDX60 selectively inhibits the translation of type II IRES-driven translation by reducing ribosome binding on type II IRES containing mRNA. In contrast, DDX60 does not reduce ribosome binding on 5’ capped mRNA. DDX60 may reduce ribosome binding to type II IRES mRNA by either inhibiting ribosome progression on the mRNA, inhibiting initiation complex assembly on the mRNA by steric hinderance, promoting detrimental structural changes in the IRES, interfering with structural rearrangements by eukaryotic initiation factors necessary for IRES function (e.g. eIF4G shown in purple, and eIF4A and 4B shown in light green), or a combination of the above.

**Figure S6.**
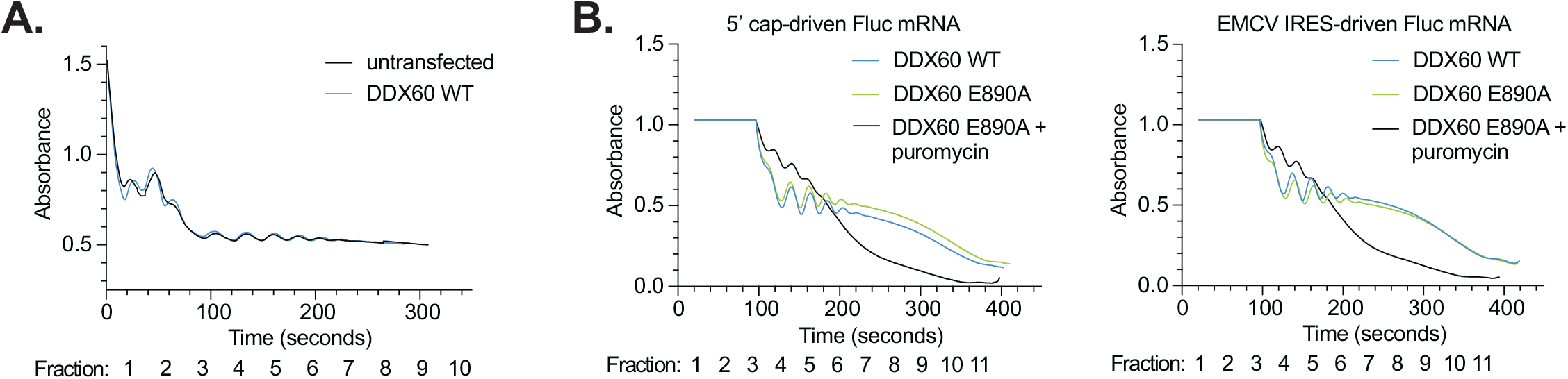
Global translation levels under DDX60 overexpression. **(A)** Polysome profile of total RNA upon DDX60 overexpression. Untransfected or DDX60 wt expressing HEK293T cells were treated with 100 μg/mL of cycloheximide for 15 minutes to arrest polysomes before being trypsinized and frozen down. All harvested cells per condition were then subjected to polysome profiling by lysing cells in the presence of cycloheximide and protease and phosphatase inhibitors, applying cell lysates to 15%-50% sucrose gradients, and subjecting to ultracentrifugation. The centrifuged gradients were run through a fractionator and total RNA in each fraction was measured by UV absorbance. Data is an average of two replicate plate of cells per condition. **(B)** Polysome profile of total RNA post transfection of DDX60 and Fluc mRNA reporters. DDX60 wt or DDX60 E890A (negative control) expressing HEK293T cells transfected with in vitro transcribed 5’ cap (left) or EMCV IRES (right) driven Fluc mRNA constructs. Duplicate samples of DDX60 E890A and 5’ cap- or EMCV IRES-driven Fluc mRNA transfected cells were treated with 200 μM puromycin as positive controls for decrease in polysomes. Cells were then treated with 100 μg/mL of cycloheximide for 15 minutes to arrest polysomes before being trypsinized and frozen down. All harvested cells per condition were then subjected to polysome profiling by lysing cells in the presence of cycloheximide and protease and phosphatase inhibitors, applying cell lysates to 15%-50% sucrose gradients, and subjecting to ultracentrifugation. The centrifuged gradients were run through a fractionator and total RNA in each fraction was measured by UV absorbance. Data is representative of 2 biological replicates from experiments conducted on separate days.

## Discussion

DDX60 is part of the superfamily-2 (SF2) DExD/H box RNA helicases that are all thought to use energy from ATP to remodel RNA structures. Structurally, these helicases contain two tandem helicase core domains with various characteristic sequence motifs, flanked by N and C-terminal extensions. They impact diverse cellular processes such as transcription, mRNA splicing, translation, and RNA turnover (Jankowsky, 2011; Pyle, 2008; Ranji & Boris-Lawrie, 2010; Sloan & Bohnsack, 2018). Given their diverse cell biological roles, DExD/H box RNA helicases are associated with the development of many diseases such as cancer, aging, neurologic and immunologic disorders, and infectious disease (Steimer & Klostermeier, 2012). *DDX60* expression has been shown to be dysregulated and associated with advanced disease and response to treatment of different cancers, such as oral cancers (Fu et al., 2016; Reyimu et al., 2021), gliomas (Zhang et al., 2021), and breast cancers (Ríos-Romero et al., 2020; Xin et al., 2020). Additionally, DNA methylation variability in *DDX60* is associated with cases of mixed connective tissue disease (Carnero-Montoro et al., 2019). Lastly, DDX60 has been observed to inhibit viral infections (Goubau et al., 2015; Ma et al., 2017; Miyashita et al., 2011; Oshiumi et al., 2015; Schoggins et al., 2011). Its mechanism of action during virus infection, however, is contested.

DDX60 was initially shown to be among the strongest inhibitors of a specific reporter HCV (Schoggins et al., 2011). Subsequent studies investigating its antiviral mechanism arrived at differing conclusions. One group suggested that DDX60 enhanced RIG-I mediated viral RNA sensing and downstream IFN signaling (Miyashita et al., 2011; Oshiumi et al., 2015), while another group suggested that DDX60 did not physically interact with RIG-I or enhance downstream IFN signaling (Goubau et al., 2015). These differences could be due to differences in knockout cells generated, IFN-driven reporters used, or other experimental differences. Here, we present an alternate but not mutually exclusive mechanism of action. We show that DDX60 is capable of specifically counteracting viral type II IRES-mediated translation while leaving host 5’ cap-mediated translation intact.

Our findings start with the unexpected observation that DDX60 can inhibit the replication of a reporter HCV containing an EMCV IRES, but not one lacking it (**Fig. 1D**). As the EMCV IRES is a commonly included element used to drive dual expression of two independent gene cassettes, it was included in the development of early reporter HCV (Date et al., 2004; During et al., 1998; Ghattas et al., 1991; Jones et al., 2010; Jones et al., 2007). However, our experiments utilizing luciferase reporters translationally regulated by different IRESs (**Fig. 2**) and comparative virology experiments utilizing different IRES carrying viruses and a chimeric virus (**Fig. 3**) combined with the differential phenotype observed with two different reporter HCV establish a strong correlation between EMCV (or type II) IRES activity and translational repression by DDX60. Additionally, our functional studies interrogating the effect of DDX60 mutations on type II IRES activity (**Fig. S2A**) strongly correlate with our bicistronic reporter HCV and EMCV IRES-driven RFP observations (**Fig. 1B, C**). Altogether, these observations warrant caution in using exogenous translational elements for reporter viruses as it may give artifactual findings when screening antiviral effectors, particularly with relation to the interferon response.

In deciphering DDX60’s mechanism of type II IRES inhibition, we posited that DDX60 may be triggering the degradation of type II IRES containing mRNAs, reducing ribosome binding on said mRNAs, acting indirectly through its previously characterized role as an enhancer of RIG-I-mediated downstream IFN activity, or a combination of the three. However, DDX60 expression did not affect the abundance of type II IRES driven mRNAs, or downstream IFN activity in concert with RIG-I (**Fig. 4 and S4**). Instead, DDX60 expression reduced accumulation of ribosomes on an active type II IRES-driven mRNA without impacting overall cellular translation or ribosome accumulation on a 5’ capped mRNA (**Fig. 5 and S6**), suggesting that DDX60 functions as a specific cell intrinsic inhibitor of protein synthesis from viral type II IRESs. How DDX60 blocks ribosome accumulation on type II IRES mRNA is still unknown (**Fig. 6**). One possibility is that DDX60 sterically hinders ribosomes from either initially binding or progressing along type II IRES mRNA. However, our RNA bindings assays (**Fig. S5**) show that DDX60 binds mRNA indiscriminately, suggesting that RNA binding alone is not determining IRES specificity. Alternatively, DDX60 may bind several different mRNAs but is displaced by translating or initiating ribosomes more efficiently on non-inhibited mRNAs. When DDX60 binds a type II IRES element, however, it may use its helicase activity to remodel the IRES, inhibiting its functions for translation initiation. Certainly, a combination of steric hinderance and IRES remodeling may also be at play. Further biochemical studies could tease apart these possibilities but would require expression and purification of the full-length ∼200 kDa DDX60 protein.

DDX60 is not the only DExD/H box RNA helicase observed to inhibit viral IRES-mediated translation. The DEAD box helicase DDX21 was recently shown to antagonize FMDV IRES, and the HCV like IRESs of classical swine fever virus and Seneca Valley virus (Abdullah et al., 2021). However, unlike DDX60, DDX21 is thought to inhibit these IRESs indirectly through upregulation of *IFN-ß* and *IL-8* mRNAs (Abdullah et al., 2021). Likewise, the ability for DDX21 to selectively inhibit IRES-mediated translation has not been tested. Other antiviral effectors that can restrict IRES-mediated translation such as PKR and RNase L dampen host mRNA translation as well (Stern-Ginossar et al., 2019). DDX60 is unique in this regard as it selectively dampens viral type II IRES-mediated translation.

Recent studies also suggest a broader role for DDX60 in translation. One study showed that kinetoplastid-DDX60 in *Trypanosoma* and *Leishmania* species associates with the 43S pre-initiation complex and possibly alters its conformation upon binding ATP to regulate translation initiation (Bochler et al., 2020). DDX60 homologs exist in every kingdom of eukaryotes (NCBI), and intriguingly, in our hands, human DDX60 can be expressed in animal cells to inhibit animal viruses (**Fig. 3**). The use of human DDX60 in hamster cells is a caveat of our FMDV and BEV-1 infection experiments, but the mammalian translation apparatus is conserved (Hernández et al., 2010), and observations from this experiment suggest a level of functional conservation for DDX60 in different species, the mammalian translation apparatus, or both. Further work comparing DDX60 sequences and functions from other species may uncover novel and/or conserved DDX60 functions. Another study found that interaction between DDX60 and the m^6^A reader protein, YTHDF1 is required for YTHDF1 to bind m^6^A modified *Traf6* (tumor necrosis factor receptor-associated factor 6) mRNA and direct its translation (Zong et al., 2021). Methylation of adenosine nucleotides at the *N*^*6*^ position (m^6^A) is the most prevalent post-transcriptional mRNA modification, generally affecting mRNA metabolism and translation (Roundtree et al., 2017; Zhao et al., 2017). Our *in vitro* transcribed mRNAs lack covalent RNA modifications which may have influenced *in vitro* DDX60-RNA interactions we observed (**Fig. S5**). Thus, future work deciphering the exact role of DDX60 in recognizing m^6^A modified *Traf6* or other m^6^A modified mRNAs may reveal how DDX60 regulates metabolism or translation of select cellularly transcribed mRNAs.

Our experiments to test how DDX60 distinguishes type II IRES containing RNAs showed that it physically binds different IRES and 5’ capped RNAs nonspecifically *in vitro* (**Fig. S5**). One possible explanation is that DDX60 has the propensity to bind many IRES types, but only interferes with specific secondary structure(s) found in type II IRESs and not the other IRES types (reviewed in (Hellen & Wimmer, 1995; Lozano & Martínez-Salas, 2015; Martinez-Salas et al., 2018). Our data demonstrate that poliovirus can be inhibited by DDX60 when its endogenous type I IRES is replaced with the type II IRES of EMCV (**Fig. 3E**), suggesting that DDX60 recognizes and/or interferes with some difference(s) between type I and type II IRESs. One possibility may be that DDX60 binds and interferes with RNA structural domains specific to type II IRESs, such as domains J and K (Hellen & Wimmer, 1995; Martinez-Salas et al., 2018). More targeted UV-crosslinking based RNA binding assays, specific deletions of domains in the EMCV IRES, or specific insertions of domains in the EMCV IRES into the poliovirus IRES can test this possibility. Alternatively, DDX60 may preferentially bind and inhibit type II IRES translation when complexed with other proteins. Indeed, other helicases have been shown to recognize their substrates by interacting with proteins via their helicase domains and N- and C-terminal extensions (Halbach et al., 2012; Lingaraju et al., 2019; Thoms et al., 2015), which we observed to both be important for DDX60 anti-IRES activity (**Fig. 1C, S2A**). An interesting possibility is that DDX60 may differentially interact and interfere with eIFs or ITAFs that are specifically involved in promoting type II IRES translation but not translation from the other IRES types (reviewed in (Hellen & Wimmer, 1995; Lozano & Martínez-Salas, 2015; Martinez-Salas et al., 2018). ITAFs differentiating type I and type II IRESs may be one group of candidates given our observations using a chimeric poliovirus (**Fig. 3E**) and that type I and type II IRESs share many eIFs (Lozano & Martínez-Salas, 2015; Steimer & Klostermeier, 2012). Immunoprecipitation and mass-spectrometry experiments may reveal proteins that interact with DDX60, and subsequent knockdown experiments may reveal the necessity of such factors for DDX60 RNA binding and/or type II IRES inhibition. Alternatively, DDX60 may interact with eIFs shared by type I and type II IRESs, but specifically interfere with the structural rearrangements made on the type II IRES by eIFs such as eIF4G and eIF4A (Kolupaeva et al., 2003). Immunoprecipitation of type I versus type II IRES bound eIFs and ITAFs could generate a list of potential proteins that contribute to DDX60 activity against type II IRESs, and *in vitro* translation, hydroxyl radical, and chemical and enzymatic assays may define possible structural changes in the type II IRES that are blocked by DDX60 but are complicated by the necessity to pull down full-length DDX60 protein in sufficient amounts.

In our analyses of different DDX60 mutants, we found that predicted ATP binding residues, helicase motif, and N and C-terminal extensions were all important for DDX60 antiviral and anti-IRES activity (**Fig. 1C, S2A**). Intriguingly, the SAT motif of DDX60 was dispensable for antiviral function (**Fig. 1C**). SAT mutants in other DEAD box helicases are able to bind RNA in an ATP dependent manner but lack RNA unwinding activity (Linder, 2006; Pause & Sonenberg, 1992; Schwer & Meszaros, 2000), suggesting that RNA binding or ATPase activity alone *may* be sufficient for DDX60’s antiviral properties. Additionally, we observed that one presumed DDX60 ATP binding/hydrolysis mutant, Q1321A, uniformly decreased both 5’ cap-driven translation and all types of IRES-driven translation (**Fig. S2A**). The analogous glutamine in the HCV encoded SF2 helicase NS3, is thought to stabilize the developing positive charge on the attacking water molecule during ATP hydrolysis (Gu & Rice, 2009). This combined with the idea that ATP hydrolysis is thought to allow progression of DExH box helicases along its RNA substrate (Steimer & Klostermeier, 2012), afford speculation that a DDX60 Q1321 mutant may bind, but not progress along an RNA substrate and allow DDX60 to have aberrant steric hinderance properties that inhibit multiple IRESs from initiating translation. Puzzlingly, we found another presumed ATP binding/hydrolysis DDX60 mutant, K791N, to uniformly increase all types of IRES-driven translation (**Fig. S2A**). The analogous lysine in NS3 is thought to help stabilize ATP binding through interaction with its ß-phosphate, and in concert with two domain VI arginine residues and a metal ion, help stabilize the developing negative charge on the ATP γ-phosphate during hydrolysis (Gu & Rice, 2009). One can speculate that perhaps a K791N mutation alters DDX60 ATP binding and hydrolysis kinetics. What these alterations are and how they contribute to regulating the activity of multiple IRESs, but not cap-mediated translation remains to be determined. Future work interrogating individual DDX60 mutants through *in vitro* ATP hydrolysis, RNA, and protein binding assays may reveal the precise enzymatic activities that DDX60 uses to enhance or inhibit different types of IRES-driven translation. As these experiments must be performed in the context of the full-length purified DDX60 protein to preserve its differential anti-IRES function (**Fig. 1B and Fig. S2A**), they will be technically challenging due to the large size and observed poor solubility of DDX60.

Lastly, IRESs are not unique to viruses and many host mRNAs have been found to initiate translation using IRES elements during conditions of stress (Jackson, 2013; Komar & Hatzoglou, 2011). Determining cellular mRNAs bound by DDX60 using an unbiased sequencing-based approach could decipher common functional or structural themes in DDX60 bound mRNAs. Likewise, a parallel sequencing-based ribosome profiling approach could reveal the compendium of mRNAs translationally regulated by DDX60.

Overall, our work uncovers DDX60 as host factor that specifically inhibits type II IRES-mediated translation, but not other IRES- or host 5’ cap-mediated translation. We show that DDX60 mechanistically acts by reducing ribosomes bound to type II IRES-driven but not 5’ cap-driven mRNAs (**Fig. 6**). DDX60 may thus work as an innate immune counter measure to specifically decrease viral protein synthesis while allowing host translation to proceed unencumbered. Future work uncovering the means for DDX60 specificity may unearth novel strategies for selective translational control in the face of a virus infection.

## Materials and Methods

### Cell lines

HEK293T (CRL-3216), A549 (CCL-185), HFF1 (SCRC-1041), and H1 clone of HeLa (CRL-1958) cells were purchased from ATCC. HEK293 ISRE reporter recombinant cell line was purchased from BPS Bioscience (Catalog #60510). HEK293T and A549 cells were cultured in DMEM (Corning™ 10013CV) supplemented with 10% FBS (Cytiva SH30071.03), penicillin and streptomycin solution (1X, Corning™ 30-002-CI), and non-essential amino acid solution (1X, Corning™ 25025CI). HFF1 cells were cultured in DMEM supplemented with 15% FBS, penicillin and streptomycin solution, and non-essential amino acid solution. HeLa and cells were cultured in MEM (Corning™ 10-009-CV) supplemented with 10% FBS and penicillin and streptomycin solution. HEK293 ISRE reporter recombinant cells were cultured in MEM supplemented with 10% FBS, penicillin and streptomycin solution, and 400 ug/mL of Geneticin™ Selective Antibiotic (G418 Sulfate) (Thermo Fisher Scientific 10131027). BHK-21 cells (baby hamster kidney cells strain 21, clone 13, ATCC CL10), obtained from the American Type Culture Collection (ATCC) were used to propagate virus stocks and to measure FMDV and BEV-1 titers. Cells were cultured in MEM supplemented with 10% FBS. BHK21/J (Leiden) 2^nd^ seed cells were cultured in DMEM (Gibco 11965118) supplemented with 10% FBS and penicillin and streptomycin solution. Huh-7 (Nakabayashi et al., 1982) and Huh-7.5 (Blight et al., 2002) were cultured in DMEM (Gibco 11965118) supplemented with 10% FBS and non-essential amino acid solution. All cell lines were tested for mycoplasma using Lookout Mycoplasma PCR kit (Sigma-Aldrich MP0035).

### Viruses

Bicistronic HCV was generated in (Schoggins et al., 2011) from Bi-YPet-Jc1FLAG2 described in (Jones et al., 2010). Monocistronic HCV J6/JFH-5AB-YPet was derived from (Horwitz et al., 2013). HCV stocks were generated by electroporation of *in vitro* transcribed RNA into Huh-7.5 cells as described in (Lindenbach et al., 2005).

FMDV was generated from the full-length serotype A12 infectious clone, pRMC35 (Rieder et al., 1993). Viruses were propagated in BHK-21 cells and were concentrated by polyethylene glycol precipitation and stored at −70 °C. Bovine enterovirus 1 (BEV-1; GenBank accession no. D00214) was obtained from Agricultural Research Service at Plum Island Animal Disease Center.

Poliovirus was rescued from pPVM-2A-144-poliovirus-GFP (GenBank accession No. pending) as described in (Burrill et al., 2013). Briefly, pPVM-2A-144-poliovirus-GFP plasmid DNA was linearized with EcoRI (NEB R3101). Purified linear DNA was then used to *in vitro* transcribe poliovirus plus stranded RNA using MEGAscript™ T7 Transcription Kit (Thermo Fisher Scientific AM1334) according to the manufacturer’s protocol. Purified RNA was then electroporated into HeLa cells. After visible cytopathic effect, media was harvested, and virus purified using ultracentrifugation.

**Table S1.**
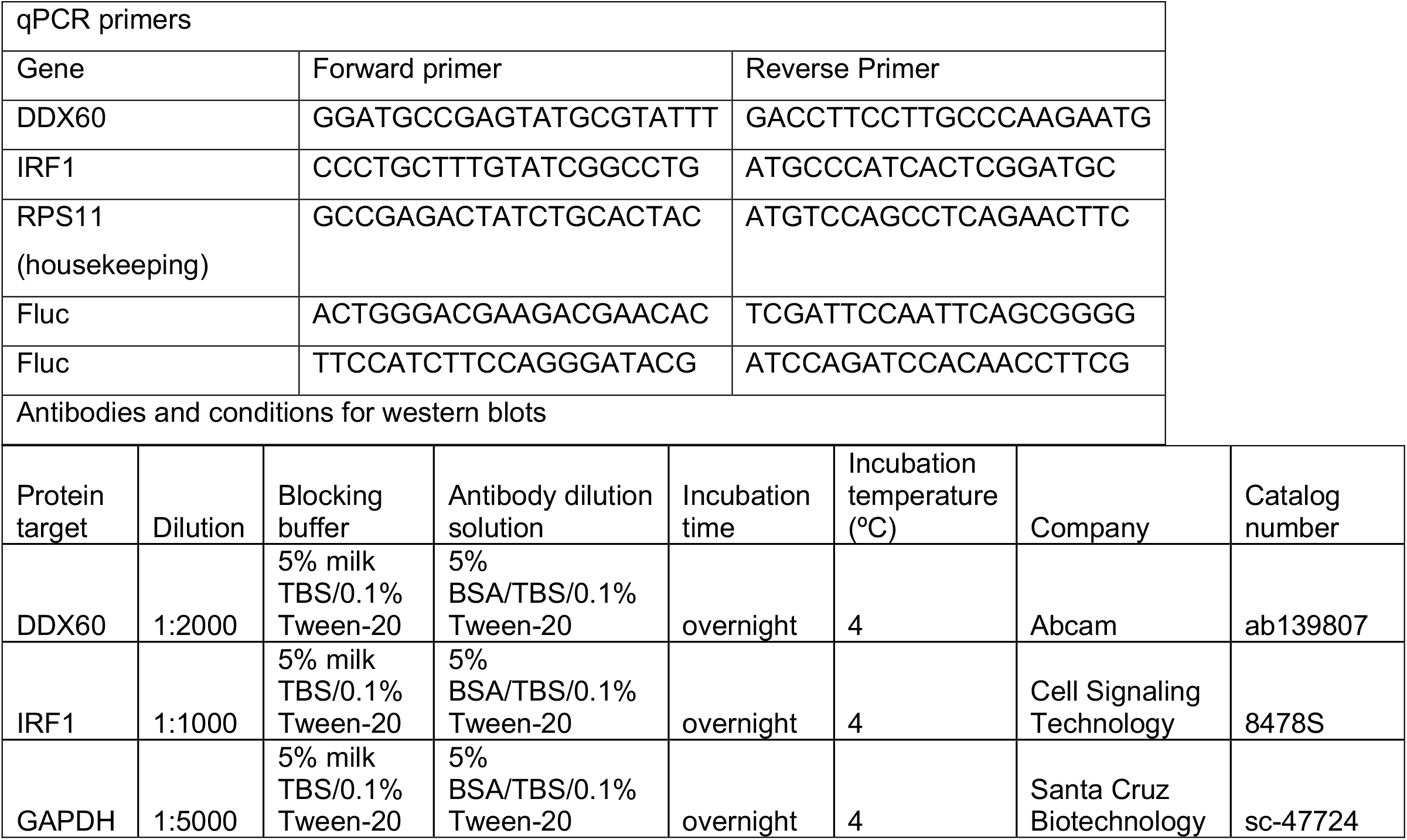
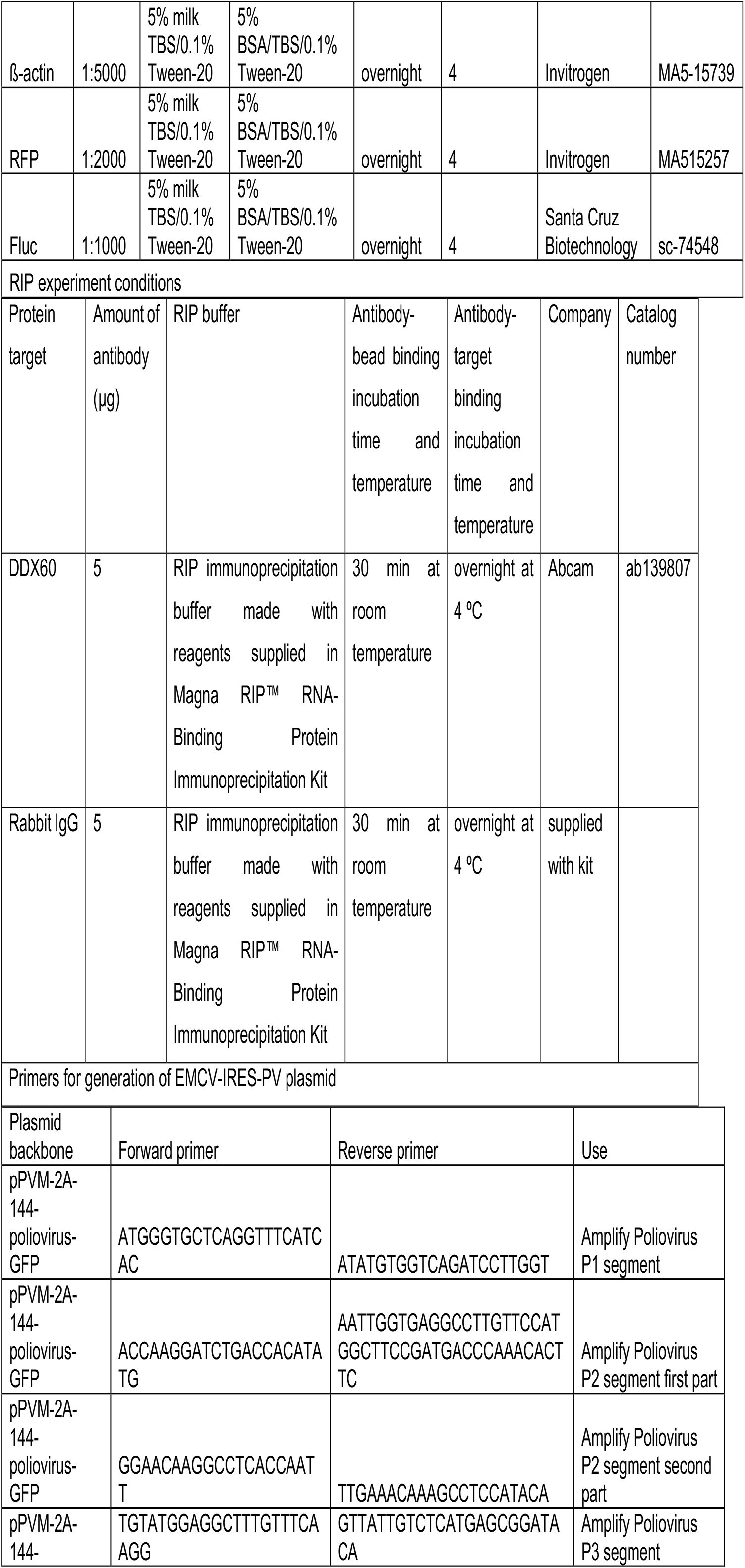
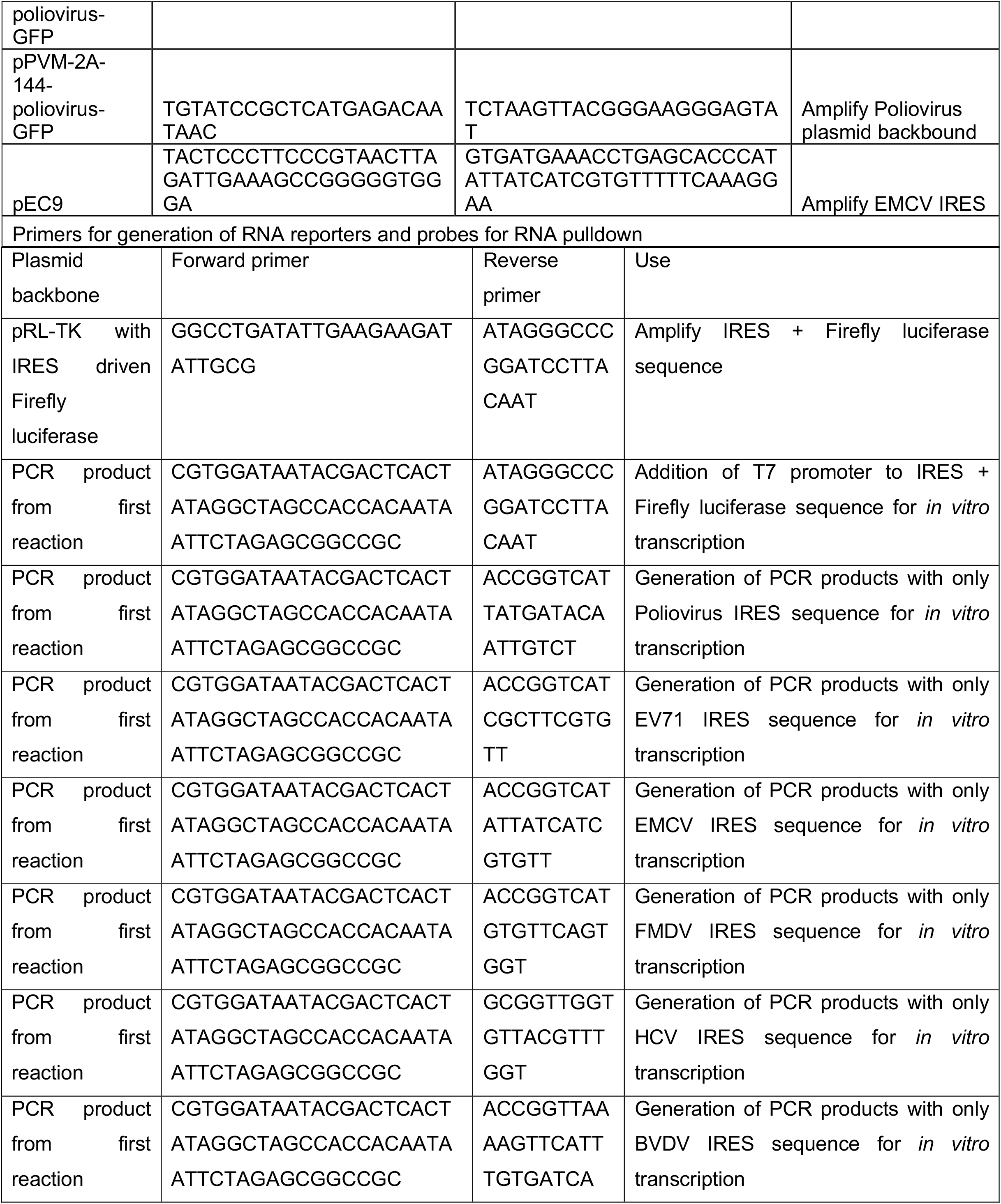
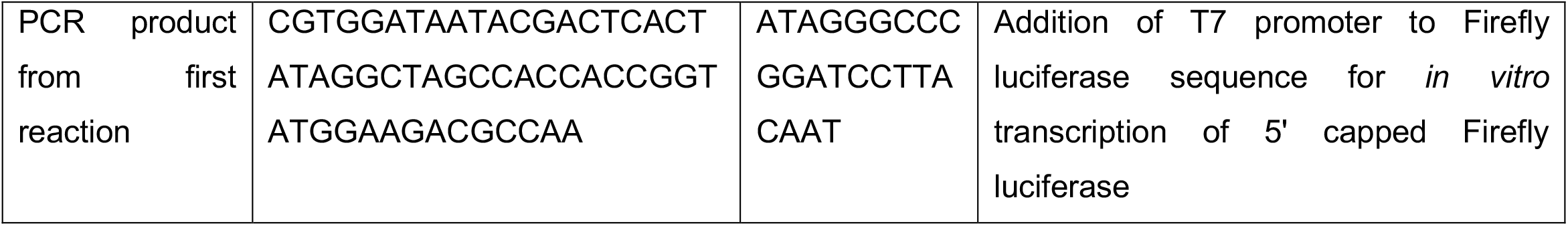

EMCV was rescued from pEC9 (a gift from Ann Palmenberg, GenBank accession No. pending). pEC9 plasmid was linearized using Sal1 (NEB R3138). Purified linear DNA was then used to *in vitro* transcribe EMCV plus stranded RNA using MEGAscript™ T7 Transcription Kit according to the manufacturer’s protocol. Purified RNA was then electroporated into HeLa cells. After visible cytopathic effect, media was harvested, and virus purified using ultracentrifugation as described in (Burrill et al., 2013).

Chimeric EMCV-IRES-PV (GenBank accession No. pending) was derived from pPVM-2A-144-poliovirus-GFP. Five PCR segments were amplified from pPVM-2A-144-poliovirus-GFP using the following primer pairs: ATGGGTGCTCAGGTTTCATCAC and ATATGTGGTCAGATCCTTGGT, ACCAAGGATCTGACCACATATG and AATTGGTGAGGCCTTGTTCCATGGCTTCCGATGACCCAAACACTTC, GGAACAAGGCCTCACCAATT and TTGAAACAAAGCCTCCATACA, TGTATGGAGGCTTTGTTTCAAGG and GTTATTGTCTCATGAGCGGATACA, TGTATCCGCTCATGAGACAATAAC and TCTAAGTTACGGGAAGGGAGTAT. EMCV IRES segment from pEC9 was amplified using primers TACTCCCTTCCCGTAACTTAGATTGAAAGCCGGGGGTGGGA and GTGATGAAACCTGAGCACCCATATTATCATCGTGTTTTTCAAAGGAA (**Table S1**). All PCR amplified segments were assembled using NEBuilder® HiFi DNA Assembly Master Mix (NEB E2621) according to the manufacturer’s protocol. Assembled plasmid was transformed into NEB Stable Competent E. Coli (NEB C3040H). Purified EMCV-IRES-PV plasmid DNA was linearized using EcoRI digestion and purified linear DNA was then used to *in vitro transcribe* EMCV-IRES-PV plus stranded RNA using MEGAscript™ T7 Transcription Kit according to the manufacturer’s protocol. Purified RNA was then electroporated into HeLa cells. After visible cytopathic effect, media was harvested, and virus purified using ultracentrifugation as described in (Burrill et al., 2013).

### Plasmids

DDX60 wild type, IRF1, and Fluc plasmid DNA was generated by (Schoggins et al., 2011) in pTRIP.CMV.IVSb.ires.TagRFP-DEST backbone (GenBank accession No. pending). Empty vector is pTRIP.CMV.IVSb.ires.TagRFP-DEST without *ccdB* suicide gene. DDX60 wild type, IRF1, and Fluc DNA was also cloned into a pTRIP.CMV.IVSb.ires.TagRFP-DEST backbone containing a puromycin selectable marker (pTRIP.CMV.IVSb.ires.TagRFP-puro-DEST). GFP plasmid is EGFP cloned into pcDNA DEST40 vector (Thermo Fisher Scientific 12274015). RIG-I plasmid DNA was generated by (Dittmann et al., 2015) in pSCRPSY lentiviral expression vector. Mutant DDX60 constructs, other ISG constructs, and bicistronic 5’ cap-driven Rluc and IRES-driven Fluc constructs (Honda et al., 2000) can be found using GenBank accession numbers: (GenBank accession No. pending).

### RNA reporters

5’ cap- and IRES-driven Fluc mRNA reporters were generated from PCR products obtained from bicistronic 5’ cap-driven Rluc and IRES-driven Fluc plasmids. First, IRES and Fluc sequences were PCR amplified from each plasmid using primers GGCCTGATATTGAAGAAGATATTGCG and ATAGGGCCCGGATCCTTACAAT. To generate IRES-driven Fluc PCR products, purified PCR product from this first PCR was used in a second round of PCR using primers CGTGGATAATACGACTCACTATAGGCTAGCCACCACAATAATTCTAGAGCGGCCGC and ATAGGGCCCGGATCCTTACAAT (**Table S1**). To generate PCR products for *in vitro* transcription of 5’ capped Fluc, PCR products from first PCR was amplified using primers CGTGGATAATACGACTCACTATAGGCTAGCCACCACCGGTATGGAAGACGCCAA and ATAGGGCCCGGATCCTTACAAT (**Table S1**). These final PCR products were purified and used for *in vitro* transcription using MEGAscript™ T7 Transcription Kit according to the manufacturer’s protocol. The *in vitro* transcription reaction for the 5’ cap driven Fluc reporter included a 1:4 ratio of Invitrogen™ Anti-Reverse Cap Analog (Fisher Scientific AM8045) and GTP solution (0.4 µl cap analog to 1.6 µl of GTP solution). After *in vitro* transcription, poly-A tails were added on using Invitrogen™ Poly(A) Tailing Kit (Thermo Fisher Scientific AM1350) according to the manufacturer’s protocol. Final reporter mRNAs were purified using Qiagen RNeasy kit (Qiagen 74106). Quality of mRNA reporters were ensured by sequencing PCR products used for *in vitro* transcription and running final RNA products on an agarose gel using NorthernMax™-Gly Sample Loading Dye (Thermo Fisher Scientific AM8551) and Invitrogen™ Ambion Millennium Markers RNA Markers Formamide (Thermo Fisher Scientific AM7151).

To generate biotin labeled and unlabeled Poliovirus IRES, EV71 IRES, EMCV IRES, FMDV IRES, HCV IRES, BVDV IRES, and 5’ capped Fluc probes, PCR products were generated from bicistronic 5’ cap-driven Rluc and IRES-driven Fluc plasmids in two rounds. First, IRES and Fluc sequences were PCR amplified from each plasmid using primers GGCCTGATATTGAAGAAGATATTGCG and ATAGGGCCCGGATCCTTACAAT (**Table S1**). Purified PCR products from this first PCR was used in a second round of PCR using primers: CGTGGATAATACGACTCACTATAGGCTAGCCACCACAATAATTCTAGAGCGGCCGC and ACCGGTCATTATGATACAATTGTCT (Poliovirus IRES), CGTGGATAATACGACTCACTATAGGCTAGCCACCACAATAATTCTAGAGCGGCCGC and ACCGGTCATCGCTTCGTGTT (EV71 IRES), CGTGGATAATACGACTCACTATAGGCTAGCCACCACAATAATTCTAGAGCGGCCGC and ACCGGTCATATTATCATCGTGTT (EMCV IRES), CGTGGATAATACGACTCACTATAGGCTAGCCACCACAATAATTCTAGAGCGGCCGC and ACCGGTCATGTGTTCAGTGGT (FMDV IRES), CGTGGATAATACGACTCACTATAGGCTAGCCACCACAATAATTCTAGAGCGGCCGC and GCGGTTGGTGTTACGTTTGGT (HCV IRES), CGTGGATAATACGACTCACTATAGGCTAGCCACCACAATAATTCTAGAGCGGCCGC and ACCGGTTAAAAGTTCATTTGTGATCA (BVDV IRES), CGTGGATAATACGACTCACTATAGGCTAGCCACCACCGGTATGGAAGACGCCAA and ATAGGGCCCGGATCCTTACAAT (5’ capped Fluc). Biotin labeled RNA probes were generated by adding 1.25 μl of 10 mM biotin-16-UTP (Millipore Sigma 11388908910) to the *in vitro* transcription reactions. Unlabeled probes were *in vitro* transcribed without biotin-16-UTP added. Final RNA was purified using Qiagen RNeasy kit. Quality of RNA was ensured by sequencing PCR products used for *in vitro* transcription and running final RNA products on an agarose gel using NorthernMax™-Gly Sample Loading Dye and Invitrogen™ Ambion Millennium Markers RNA Markers Formamide.

### Interferon-ß treatment time course experiments

HEK293T, A549, HFF1, or HeLa cells were plated at a density of 5E4 (A549 and HFF1) or 1E5 (HEK293T and HeLa) cells/cm^2^ in a 24-well plate with 2 wells for 0.1% BSA/PBS treatment and 2 wells for 500 U/mL IFN-ß (dissolved in 0.1% BSA/PBS, Millipore Sigma IF104) treatment. Wells with HEK293T cells were coated with poly-L-ornithine (Sigma-Aldrich P2533) prior to plating. Each cell line was treated with either 0.1% BSA/PBS or 500 U/mL IFN-ß. 0-, 6-, 12-, 24-, or 48-hours post treatment, cells were washed 1x with PBS (Fisher Scientific MT21031CV) and lysed in either 350 µl buffer RLT (supplied with Qiagen 74106) supplemented with 2-mercaptoethanol at a 1:100 dilution (Sigma-Aldrich M3148) or 200 µl LDS sample buffer (diluted to 1X with molecular biology grade water, Invitrogen™ B0007) supplemented with cOmplete™, Mini, EDTA-free Protease Inhibitor Cocktail (Roche 11836170001) and Pierce™ Phosphatase Inhibitor Mini Tablets (Thermo Fisher Scientific A32957). RNA was isolated using Qiagen RNeasy kit (Qiagen 74106) and cDNA synthesized using SuperScript™ III First-Strand Synthesis System (Thermo Fisher Scientific 18080051). RT-qPCR was performed using Applied Biosystems™ PowerUp™ SYBR™ Green Master Mix (Applied Biosystems™ A25918), Applied Biosystems™ QuantStudio 3 real-time PCR system, and primers for genes listed in **Table S1** (20 ul reaction volume per well consisting of 3 µl 3 µM primer mix, 10 µl PowerUp™ SYBR™ Green Master Mix, 5 µl cDNA, and 2 µl RNase-free water). Data was analyzed using delta-delta Ct method devised by (Livak & Schmittgen, 2001) after normalizing Ct values to housekeeping gene RPS11. Western blot was performed by boiling samples at 95 °C for 5 minutes and loading samples on a NuPAGE™ 3 to 8%, Tris-Acetate, 1.5 mm, gel. Proteins were then transferred onto a PVDF membrane using iBlot 2 transfer system (Thermo Fisher Scientific IB24001). Membrane was blocked in 5% Thermo Scientific™ Oxoid™ Skim Milk Powder (Fisher Scientific OXLP0031B) TBS/0.1% Tween-20 solution (Fisher Scientific BP337-500). After 3×10 minute washes with TBS/0.1% Tween-20, membrane was probed for antibodies of interest using the conditions listed in **Table S1**. After primary antibody incubation, membrane was washed with TBS/0.1% Tween-20 for 3×10 minutes and probed with LI-COR IRDye® 800CW Donkey anti-Rabbit IgG and IRDye 680RD Donkey Anti-Mouse IgG secondary antibodies for 1 hour at room temperature. After 3×10 minute washes with TBS/0.1% Tween-20, proteins were visualized using Odyssey® DLx Imaging System. Band intensities were measured using ImageJ2 (2.3.0/1.53f). Region of interests (ROI) were defined for loading control and protein of interest (POI) bands and mean gray values for each band was measured. Background mean gray values using the defined ROIs were also measured from parts of each lane without any protein bands. Inverted pixel densities were calculated by subtracting the mean gray values from 255. Net densities for loading control and POI bands were calculated by subtracting the inverted pixel densities of the background values from the inverted pixel densities of either the loading control or POI bands. Net densities of POI bands were then divided by the net densities of loading control bands to define the final relative quantification values.

### Western blots to confirm expression of transgenes

HEK293T cells were plated at a density of 1E5 cells/cm^2^ on a 24-well poly-L-ornithine coated plate. The next day, cells were transfected with 1 µg of DDX60 wt or mutant constructs using TransIT®-LT1 Transfection Reagent (Mirus Bio MIR 2300) at a 1:3 (µg to µl) DNA to lipid reagent ratio in a total volume of 50 µl of Opti-MEM (Thermo Fisher Scientific 51985034) according to the TransIT®-LT1 manufacturer’s protocol for 24-well plate. 16-18 hours post transfection, media on cells were changed to DMEM supplemented with 1.5% FBS, penicillin and streptomycin, and non-essential amino acids. 48 hours post transfection, cells were washed 1x with PBS and lysed in 200 µl LDS sample buffer (diluted to 1X with molecular biology grade water) supplemented with cOmplete™, Mini, EDTA-free Protease Inhibitor Cocktail and Pierce™ Phosphatase Inhibitor Mini Tablets. Western blot was performed by boiling samples at 95 °C for 5 minutes and loading samples on a NuPAGE™ 3 to 8%, Tris-Acetate, 1.5 mm, gel. Proteins were then transferred onto a PVDF membrane using iBlot 2 transfer system. Membrane was blocked in 5% Oxoid™ Skim Milk Powder TBS/0.1% Tween-20 solution. After 3×10 minute washes with TBS/0.1% Tween-20, membrane was probed for antibodies of interest using the conditions listed in **Table S1**. After primary antibody incubation, membrane was washed with TBS/0.1% Tween-20 for 3×10 minutes and probed with LI-COR IRDye® 800CW Donkey anti-Rabbit IgG and IRDye 680RD Donkey Anti-Mouse IgG secondary antibodies for 1 hour at room temperature. After 3×10 minute washes with TBS/0.1% Tween-20, proteins were visualized using Odyssey® DLx Imaging System. Band intensities were measured using ImageJ2 as described above.

### Antiviral assays with HCV reporters

HCV antiviral assays were conducted as described in (Schoggins et al., 2011). Huh-7 cells were plated on 24-well plates at a density of 7E4 cells/well. The next day, growth medium was changed to DMEM supplemented with 1.5% FBS, penicillin and streptomycin, non-essential amino acids. 400 ng DNA/well of Fluc, IRF1, DDX60 wt, or DDX60 mutant constructs were transfected using Lipofectamine 2000 (Thermo Fisher Scientific 11668019) in a total volume of 1 mL/well. Plates were centrifuged at 1000g for 30 min at 37 °C. 5 hours later, the medium was changed to DMEM supplemented with 10% FBS, penicillin and streptomycin, non-essential amino acids. 48 hours post transfection, cells were washed 1x with PBS and infected with HCV-Ypet bicistronic or monocistronic HCV J6/JFH-5AB-YPet reporter at a dose yielding approximately 50% infected cells as determined by flow-cytometry based infectivity assays on Fluc transfected cells. 72 hpi, adherent cells were collected into 200 µl Accumax Cell Aggregate Dissociation Medium (Thermo Fisher Scientific 00-4666-56) and transferred to a 96-well plate. Cells were pelleted at 1000g for 5 min at 4 °C and resuspended in 1% paraformaldehyde fixation solution for 1 h. Cells were pelleted once again by centrifugation at 1,000g for 5 min at 4 °C and resuspended in cold PBS supplemented with 3% FBS and stored at 4 °C until flow cytometry analysis. Samples were analyzed in a 96-well-based high-throughput manner using an LSRII-HTS flow cytometer (BD Biosciences). Data were analyzed using FlowJo software with a 0.1% compensation matrix. Percent of Ypet expressing cells in RFP-positive cells was determined and plotted.

### In cell plasmid-based reporter assays

HEK293T cells were plated on a poly-L-ornithine coated 48-well plate at a density of 6E4 cells/cm^2^. The next day, cells were co-transfected with 200 ng of bicistronic 5’ cap driven Rluc and IRES driven Fluc plasmid and 300 ng of GFP alone, or 290 ng GFP + 10 ng IRF1, or 200 ng GFP + 100 ng IRF1, or 300 ng IRF1 alone, or 290 ng GFP + 10 ng DDX60 wt, or 200 ng GFP + 100 ng DDX60 wt, or 300 ng DDX60 wt alone, or 290 ng GFP + 10 ng DDX60 E890A, or 200 ng GFP + 100 ng DDX60 E890A, or 300 ng DDX60 E890A alone. Transfections were performed using X-tremeGENE 9 DNA transfection reagent (Roche), with 1.2 µl X-tremeGENE reagent per 500 µg DNA reaction. 16-18 hours post transfection, media on cells were changed to DMEM supplemented with 1.5% FBS, penicillin and streptomycin, and non-essential amino acids. 48 hours post transfection, cells were lysed using 1X passive lysis buffer (5X passive lysis buffer diluted in PBS) supplied in Promega Dual-Luciferase™ Reporter assay system (Fisher Scientific PR-E1980) according to the manufacturer’s protocol with the addition of 1 freeze-thaw cycle step, freezing for at least 1 hour at -80 °C before thaw. Renilla and Firefly luciferase activity was then assayed using Promega Dual-Luciferase™ Reporter assay system according to the manufacturer’s protocol.

### In cell RNA-based reporter assays

HEK293T cells were plated on a poly-L-ornithine coated 48-well plate at a density of 3E4 cells/cm^2^. The next day, cells were transfected with either 500 ng GFP, IRF1, DDX60 wt, or DDX60 mutant constructs using TransIT®-LT1 Transfection Reagent at a 1:3 (µg to µl) DNA to lipid reagent ratio in a total volume of 30 µl of Opti-MEM according to the TransIT®-LT1 manufacturer’s protocol for a 48-well plate. 16-18 hours post transfection, media on cells were changed to DMEM supplemented with 1.5% FBS, penicillin and streptomycin, and non-essential amino acids. 48 hours post transfection, cells were transfected with 0.2-0.3 pmol of either 5’ cap-, Poliovirus IRES-, EV71 IRES-, EMCV IRES-, FMDV IRES-, HCV IRES-, or BVDV IRES-driven Fluc mRNA reporters using TransIT®-mRNA Transfection Kit (Mirus Bio MIR 2250) at a 1:2 (µg to µl) RNA to lipid reagent ratio in a total volume of 30 µl of Opti-MEM according to the TransIT®-mRNA manufacturer’s protocol for a 48-well plate. 24 hours post mRNA transfection, cells were lysed using 1X passive lysis buffer (5X passive lysis buffer diluted in PBS) Passive Lysis Buffer (Promega E1941) according to the manufacturer’s protocol with the addition of 1 freeze-thaw cycle step, freezing for at least 1 hour at -80 °C before thaw. Firefly luciferase activity was then assayed using Promega Luciferase Assay System (E4550) according to the manufacturer’s protocol. Luminescence activity was measured using BioTek Synergy™ HTX Multi-Mode Microplate Reader with Agilent BioTek Dual Reagent Injector system.

### Generation of stable HeLa cell lines using lentivirus transduction

Lentiviral pseudoparticles generation was carried out as in (Schoggins et al., 2011). 1E6 HEK293T cells in 6-well plates were co-transfected with plasmids expressing the Fluc, IRF1, DDX60 wt, or DDX60 E890A pTRIP.CMV.IVSb.ISG.ires.TagRFP.puro proviral DNA, HIV-1 gag–pol and VSV-G in a ratio of 1/0.8/0.2, respectively. For each transfection, 6 µl TransIT®-LT1 Transfection Reagent was combined with 2 µg total DNA in a total volume of 200 µl Opti-MEM, allowed to mix for 15 minutes, and added dropwise to the 6-well plate. Transfections were carried out for 16-18 hours, followed by a medium change to DMEM supplemented with 1.5 % FBS, penicillin and streptomycin, and non-essential amino acids. Supernatants were collected at 48h and 72h, pooled, cleared by centrifugation and passing through a 0.45 µm filter, and stored at -80 °C.

HeLa cells were plated on a 6-well plate at 1E6 cells/well. The next day, cells were transduced with undiluted (DDX60 wt and DDX60 E890A) or 1:50 diluted (Fluc and IRF1) lentiviral pseudoparticles by spinoculation at 1250g for 45 minutes at 37 °C in DMEM supplemented with 1.5% FBS, 20 mM HEPES (Gibco™ 15630080), and 4 µg/mL polybrene (Sigma-Aldrich TR-1003-G). Undiluted DDX60 wt and E890A lentiviral pseudoparticles had to be used due to lower viral titers. 5 hours post spinoculation, cell medium was changed to MEM supplemented with 1.5% FBS and penicillin and streptomycin. 48 hours post transduction, cells were trypsinized and plated in T175 flasks in MEM supplemented with 1.5% FBS, penicillin and streptomycin, and 2 µg/mL puromycin (Sigma-Aldrich P8833). Medium was replaced on cells every 48 hours until cells were ∼90% confluent. Afterward, cells were trypsinzed and frozen down in MEM supplemented with 10% FBS, penicillin and streptomycin, and 5% DMSO.

### Antiviral assays with IRES containing viruses

HeLa cells stably expressing Fluc, IRF1, DDX60 wt, or DDX60 E890A were thawed from frozen vials and passaged once under 2 µg/mL puromycin selection. Cells were then plated on a 6-well plate at a density of 2E4 cells/cm^2^ in MEM supplemented with 10% FBS, penicillin and streptomycin, and 2 µg/mL puromycin. When plating cells, approximately 5E5 of the remaining cells were lysed for western blot analysis to check for transgene expression (see methods above for collection and western blot details). An additional well was used for seeding untransduced HeLa cells at the same density in MEM supplemented with 10% FBS and penicillin and streptomycin. Two days later, untransduced HeLa cells were trypsinized and counted. Transduced HeLa cells were washed once with PBS + Ca and Mg (Corning™ 21030CV) supplemented with 0.3% BSA (Equitech-Bio BL62-0500) and subsequently infected with EMCV, poliovirus, or EMCV-IRES-PV at an MOI of 0.001 based on the untransduced HeLa cell count numbers in a volume of 500 µl of PBS + Ca and Mg supplemented with 0.3% BSA. Approximately 50 µl of input virus was kept frozen at -80 °C for titration. This is considered the 0h time point. Infection was carried out for 1 hour at 37 °C with gentle shaking. After 1 hour, virus was aspirated, and cells washed 3x with 1 mL of PBS + Ca and Mg supplemented with 0.3% BSA. After the final wash, medium was changed to 2 mL of MEM supplemented with 1.5% FBS and penicillin and streptomycin. 120 µl of cell medium was taken for each time point and frozen at -80 °C. Cell medium was replaced with 120 µl of fresh MEM supplemented with 1.5% FBS and penicillin and streptomycin after each time point to preserve the final volume of cell medium. After the experiment, all equipment and waste were disinfected with 10% bleach.

Virus collected for each time point was titered using plaque assay. For this, HeLa cells were plated on 12-well plates at 3.5E5 cells/well. The next day, virus time points to be titered were serially diluted from 1E-1 to 1E-6 for time points <24h and 1E-3 to 1E-8 for time points >24h in a final volume of 250 µl of PBS + Ca and Mg supplemented with 0.3% BSA. Cells plated on 12-well plates were washed once with PBS + Ca and Mg supplemented with 0.3% BSA and subsequently infected with 200 µl of serially diluted virus. Infection was carried out for 1 hour at 37 °C with gentle shaking. After 1 hour, virus was aspirated, cells were washed 1x with PBS + Ca and Mg supplemented with 0.3% BSA, and medium was changed two 1 mL of overlay medium. Overlay medium for 2×12 well plates: 12.5 mL 2X DMEM (Gibco™ 12100046), 8.3 mL 4% Avicel, 250 µl DEAE Dextran (Sigma-Aldrich 30461), 250 µl Glutamax (Thermo Fisher Scientific 35050061), 250 µl non-essential amino acid solution, 250 µl penicillin and streptomycin, 1.23 mL 7.5% sodium bicarbonate (Gibco™ 25080094), and 2 mL molecular biology grade water (Fisher Scientific MT46000CM). Cells were incubated in a 37 °C incubator 18 – 24 hours for EMCV time points and 40 – 48 hours for poliovirus and EMCV-IRES-PV time points. To stop plaque assays and count viral plaques, cells were fixed by adding 1 mL of 2% Formaldehyde (Fisher Scientific BP531-500) onto the overlay medium for at least 15 minutes at room temperature. Overlay medium was then aspirated, cells washed 1x with PBS, and stained with 400 µl of crystal violet working solution for at least 15 minutes at room temperature. Crystal violet working solution consists of 40 mL 1% crystal violet stock solution, 80 mL methanol, and 300 mL water. 1% Crystal violet stock solution consists of 10 g crystal violet, 200 mL ethanol, and 800 mL water. After crystal violet staining, crystal violet working solution was removed (can be reused) and plates plunged into a bucket of water. Plaques were counted for a well with 5 – 20 plaques and viral titers calculated by using the following formula: Titer (PFU/mL) = well plaque count / (0.200 mL x dilution factor).

Antiviral assays using FMDV, and BEV-1 were performed in a biosafety level 3 facility at the Plum Island Animal Disease Center. BHK-J cells stably expressing empty vector, IRF1, DDX60 wt, or DDX60 E890A were infected with BEV-1 or FMDV at MOI 1. After one hour of adsorption, cells were rinsed once with 150 mM NaCl, 20 mM morpholineethanesulfonic acid (MES; pH 6) to inactivate unadsorbed virus and once with MEM to neutralize the MES, followed by addition of MEM and incubation at 37°C. Supernatants and cell lysates were collected at 5 hpi and titers were determined via plaque assay on BHK-21 cells.

### RNA abundance assay using IRES driven Fluc reporter RNAs

HEK293T cells were plated on a poly-L-ornithine coated 48-well plates at a density of 3E4 cells/cm^2^ (at least 2 wells per condition). The next day, cells were transfected with either 500 ng of DDX60 wt or DDX60 E890A constructs using TransIT®-LT1 Transfection Reagent at a 1:3 (µg to µl) DNA to lipid reagent ratio in a total volume of 30 µl of Opti-MEM according to the TransIT®-LT1 manufacturer’s protocol for a 48-well plate. 16-18 hours post transfection, media on cells were changed to DMEM supplemented with 1.5% FBS, penicillin and streptomycin, and non-essential amino acids. 48 hours post transfection, cells were transfected with 0.2-0.3 pmol of either 5’ cap-, Poliovirus IRES-, EV71 IRES-, EMCV IRES-, FMDV IRES-, HCV IRES-, or BVDV IRES-driven Fluc mRNA reporters using TransIT®-mRNA Transfection Kit at a 1:2 (µg to µl) RNA to lipid reagent ratio in a total volume of 30 µl of Opti-MEM according to the TransIT®-mRNA manufacturer’s protocol for a 48-well plate. 24 hours post mRNA transfection, cells were trypsinzed and half of the cell volume was lysed using 1X passive lysis buffer (5X passive lysis buffer diluted in PBS) Passive Lysis Buffer according to the manufacturer’s protocol with the addition of 1 freeze-thaw cycle step, freezing for at least 1 hour at -80 °C before thaw. Firefly luciferase activity was then assayed using Promega Luciferase Assay System according to the manufacturer’s protocol. Luminescence activity was measured using BioTek Synergy™ HTX Multi-Mode Microplate Reader with Agilent BioTek Dual Reagent Injector system. The other half of the cell volume was lysed using 350 µl buffer RLT (supplied with Qiagen 74106) supplemented with 2-mercaptoethanol at a 1:100 dilution. RNA was isolated using Qiagen RNeasy kit and cDNA synthesized using SuperScript™ III First-Strand Synthesis System. RT-qPCR was performed using Applied Biosystems™ PowerUp™ SYBR™ Green Master Mix, Applied Biosystems™ QuantStudio 3 real-time PCR system, and primers for genes listed in **Table S1** (20 ul reaction volume per well consisting of 3 µl 3 µM primer mix, 10 µl PowerUp™ SYBR™ Green Master Mix, 5 µl cDNA, and 2 µl RNase-free water). Data was analyzed using delta-delta Ct method devised by (Livak & Schmittgen, 2001) after normalizing Ct values to housekeeping gene RPS11.

### ISRE reporter assays

HEK293 ISRE reporter recombinant cells were plated on a poly-L-ornithine coated 48-well plates at a density of 6E4 cells/cm^2^ (at least 2 wells per condition). The next day, cells were transfected with either: 100 ng RIG-I + 400 ng of GFP, 125 ng RIG-I + 375 ng GFP, 150 ng RIG-I + 350 ng GFP, 175 ng RIG-I + 325 ng GFP, 100 ng RIG-I + 100 ng DDX60 wt + 300 ng GFP, 125 ng RIG-I + 125 ng DDX60 wt + 250 ng GFP, 150 ng RIG-I + 150 ng DDX60 wt + 200 ng GFP, 175 ng RIG-I + 175 ng DDX60 wt + 150 ng GFP, 100 ng RIG-I + 100 ng DDX60 E890A + 300 ng GFP, 125 ng RIG-I + 125 ng DDX60 E890A + 250 ng GFP, 150 ng RIG-I + 150 ng DDX60 E890A + 200 ng GFP, or 175 ng RIG-I + 175 ng DDX60 E890A + 150 ng GFP. Transfections were carried out using TransIT®-LT1 Transfection Reagent at a 1:3 (µg to µl) DNA to lipid reagent ratio in a total volume of 30 µl of Opti-MEM according to the TransIT®-LT1 manufacturer’s protocol for a 48-well plate. 16-18 hours post transfection, media on cells were changed to MEM supplemented with 10% FBS, penicillin and streptomycin, and 400 ug/mL geneticin. Two days post transfection, cells were either treated with PBS or were transfected with 0.641 pmol (∼250 ng) of low molecular weight poly(I:C) (Invivogen tlrl-picw) using Lipofectamine™ RNAiMAX Transfection Reagent (Thermo Fisher Scientific 13778075) in a final volume of 30 µl of Opti-MEM according to RNAiMAX manufacturer’s protocol. One well of untransfected cells was also treated with 500 U/mL of IFN-ß for 16-18 hours. 24 hours post RNA transfection, one well per condition of cells were lysed using 1X passive lysis buffer (5X passive lysis buffer diluted in PBS) Passive Lysis Buffer according to the manufacturer’s protocol with the addition of 1 freeze-thaw cycle step, freezing for at least 1 hour at -80 °C before thaw. Firefly luciferase activity was then assayed using Promega Luciferase Assay System according to the manufacturer’s protocol. Luminescence activity was measured using BioTek Synergy™ HTX Multi-Mode Microplate Reader with Agilent BioTek Dual Reagent Injector system. Another well per condition of cells were lysed in 100 µl LDS sample buffer (diluted to 1X with molecular biology grade water) supplemented with cOmplete™, Mini, EDTA-free Protease Inhibitor Cocktail and Pierce™ Phosphatase Inhibitor Mini Tablets. Western blots were run and analyzed for DDX60, RIG-I, ß-actin, and GAPDH expression as described in western blot methods described above.

### Biotin labeled RNA pulldown

Biotin labeled RNA pulldown experiments were adapted from (Hung et al., 2016). HEK293T cells were plated on a poly-L-ornithine coated 10-cm dish at a density of 4E4 cells/cm^2^. The next day, cells were transfected with 15 µg of DDX60 wt construct using TransIT®-LT1 Transfection Reagent at a 1:3 (µg to µl) DNA to lipid reagent ratio in a total volume of 1.5 mL of Opti-MEM according to the TransIT®-LT1 manufacturer’s protocol for a 10-cm dish. 16-18 hours post transfection, medium on cells was changed to DMEM supplemented with 1.5% FBS, penicillin and streptomycin, and non-essential amino acids. 48 hours post transfection, cells were washed with 2 mL of PBS and subsequently lysed at 4 °C with 300 µl 3-[(3-cholamidopropyl)-dimethylammonio]-1-propanesulfonate (CHAPS) lysis buffer (10 mM Tris-HCl [pH 7.4], 1 mM MgCl2, 1 mM EGTA, 0.5% CHAPS, 10% glycerol, 0.1 mM phenyl-methylsulfonyl fluoride [PMSF], 5 mM β-mercaptoethanol) with gentle shaking for 30 min. Cell lysates were then transferred to a 1.5 mL tube and centrifuged at 10000g for 10 min at 4 °C. Supernatants were then transferred to new tubes. Protein collected was quantified using Pierce™ BCA Protein Assay Kit (Thermo Fisher Scientific 23225) according to the manufacturer’s protocol. Samples were then frozen at -80 °C.

Before RNA pulldown, cell lysates were pre-cleared to remove proteins that have a background binding to streptavidin MagneSphere paramagnetic particles. First, streptavidin MagneSphere paramagnetic particles (Promega, Z5481) were washed 3 times with 1 mL of RNA mobility buffer without heparin (5 mM HEPES pH 5.5, 40 mM KCl, 0.1 mM EDTA, 2 mM MgCl_2_, 2 mM DTT, 1 U RNasin). To wash, place tubes containing streptavidin MagneSphere paramagnetic particles on a magnetic stand (DynaMag™-2 Magnet, Life Technologies 12321D) for 30s – 1 min, remove supernatant, and add 1 mL of RNA mobility buffer without heparin to each tube and invert tubes up and down to resuspend paramagnetic beads. Repeat the mentioned steps for 3 washes, removing the supernatant on the last wash, but not adding back any RNA mobility buffer. Cell lysates were then pre-cleared by mixing appropriate volume of cell lysate for the number of pulldowns (200 µg protein per pulldown) with streptavidin MagneSphere paramagnetic particles and incubating at room temperature for 10 min on a rocking nutator. The mixture was then placed on a magnetic stand for 30s – 1 min and the supernatant was transferred to a new sterile 1.5 mL tube on ice. The remaining streptavidin MagneSphere paramagnetic particles were resuspended in 25 µl of 2X SDS sample buffer, boiled at 95 °C for 10 min, and frozen at -80 °C.

For RNA pulldown, 200 ug of DDX60 expressing cell lysate was well mixed with 12.5 pmol of unlabeled or biotin labeled Polio IRES, EMCV IRES, HCV IRES, or 5’ capped Firefly luciferase RNA in RNA mobility buffer with heparin (5 mM HEPES pH 5.5, 40 mM KCl, 0.1 mM EDTA, 2 mM MgCl_2_, 2 mM DTT, 1 U RNasin, 0.25 mg/mL Heparin). 1% of cell extract was kept for an input sample. The cell lysate and RNA mixture was incubated at 30 degrees for 15 minutes with gentle shaking. In the meantime, streptavidin MagneSphere paramagnetic particles were washed 3 times with 1 mL of RNA mobility buffer without heparin (split 1 tube of streptavidin MagneSphere paramagnetic particles for every 2 RNA pulldowns). After the last wash, streptavidin MagneSphere paramagnetic particles were combined and resuspended in 400 µl of RNA mobility buffer without heparin per RNA pulldown. 400 µl of resuspended streptavidin MagneSphere paramagnetic particles were then added to cell lysate and RNA mixture and incubated for 10 min at room temperature rocking on a nutator. Pulled-down complexes were then washed 5 times with RNA mobility buffer without heparin. After the last wash, streptavidin MagneSphere paramagnetic particles with bound RNA and protein were resuspended in 25 µl of 2X SDS sample buffer. 1 volume of 2X SDS sample buffer was also added to 1 volume of input sample. All samples were boiled at 95 °C for 10 min, and frozen at -80 °C. To analyze bound proteins, samples were spun down at maximum speed for 1 min and supernatants used for western blots (see western blot protocol above for details).

### RNA immunoprecipitation assay

RNA immunoprecipitation (RIP) assays were performed using Magna RIP™ RNA-Binding Protein Immunoprecipitation Kit (EMD Millipore 17-700). HEK293T cells were plated on 15-cm dishes coated with poly-L-ornithine at a density of 4E4 cells/cm^2^. The next day, cells were either left untransfected or transfected with 70 µg of DDX60 wt construct using TransIT®-LT1 Transfection Reagent at a 1:3 (µg to µl) DNA to lipid reagent ratio in a total volume of 3.5 mL of Opti-MEM according to the TransIT®-LT1 manufacturer’s protocol. 16-18 hours post transfection, cell media on both untransfected and transfected cells were changed to DMEM supplemented with 1.5% FBS, penicillin and streptomycin, and non-essential amino acids. 48 hours post transfection, cells were transfected with ∼30 – 40 pmol of either 5’ cap or EMCV IRES driven Fluc reporter RNA using TransIT®-mRNA Transfection Kit at a 1:2 (µg to µl) RNA to lipid reagent ratio in a total volume of 3.5 mL of Opti-MEM according to the TransIT®-mRNA manufacturer’s protocol. The next day, cells were lysed for RIP lysate preparation using the Magna RIP™ RNA-Binding Protein Immunoprecipitation Kit and protocol. Briefly, 115 µl of RIP lysis buffer with added protease inhibitor (0.5 µl) and RNase inhibitor cocktail (0.25 µl) was prepared for each sample. Cells on the plate were washed twice with 10 mL of ice-cold PBS. Then, cells were scraped off the plates in 10 mL of ice-cold PBS per plate and transferred to 15 mL tubes. Cells were collected by centrifugation for 5 minutes at 1500 rpm at 4 °C. The supernatant was then discarded, and cell pellet was resuspended in an equal volume of RIP lysis buffer and mixed until the mixture appeared homogenous. Lysate was then incubated on ice for 5 minutes. Cell lysate was dispensed into 1.5 mL tubes in 200 µl aliquots. 100 µl of cell lysate is used per RIP and a positive and negative antibody is used for each experiment making 200 µl a single use aliquot. All aliquots were frozen at -80 °C.

On the day of RIP experiment, the appropriate number of tubes per RIP was prepared and 50 µl of magnetic beads placed into each tube. Magnetic beads were washed twice with 0.5 mL of RIP wash buffer. To wash, magnetic beads were placed on a magnetic stand for 30s – 1 min, supernatant discarded, 0.5 mL of RIP wash buffer added, and tubes vortexed briefly (setting 7). After the final wash, magnetic beads in each tube were resuspended in 100 µl of RIP wash buffer. Next, the appropriate about of antibody (5 µg in this case) was added to each tube and antibody-bead complexes were allowed to form by incubating all tubes on a rotating nutator at room temperature for 30 min (**Table S1**). Tubes were then centrifuged briefly and placed on a magnetic stand, supernatant removed, and beads washed twice with RIP wash buffer. After the last wash, 0.5 mL of RIP wash buffer was added to each tube and vortexed briefly (setting 7) before incubating tubes on ice. Next, enough RIP immunoprecipitation buffer was prepared for the appropriate number of RIPs according to Magna RIP™ RNA-Binding Protein Immunoprecipitation Kit protocol. 900 µl of RIP immunoprecipitation buffer was added to each tube on ice. RIP lysates were then thawed quickly and centrifuged at 14000 rpm for 10 minutes at 4 °C. 10 µl of supernatant was used for input samples (one for western blot and one for RNA preparation) and placed on ice. 100 µl of supernatant was added to the appropriate antibody-bead complexes. All immunoprecipitation tubes were then left rotating overnight on a nutator at 4 °C. An equal volume of 2X SDS was added to western blot input samples, samples were boiled at 95 °C for 5 minutes, and then stored at -80 °C. The RNA input sample was simply frozen at – 80 °C.

The next day, immunoprecipitation tubes were centrifuged briefly and placed on a magnetic stand. 100 µl of the supernatant was saved as unbound protein fraction for western blot. 100 µl of 2X SDS was added to the to the supernatant collected for western blot, boiled at 95 °C for 5 minutes, and stored at -80 °C. The rest of the supernatant was discarded. Beads were then washed a total of 6 times with 0.5 mL of cold RIP wash buffer. 50 µl out of 500 ul of the bead suspension during the last wash was saved from each tube to test the efficiency of immunoprecipitation by western blot. Proteins were eluted off the 50 µl of beads saved by resuspending the beads in 20 µl of 1X SDS sample buffer following by heating at 95 °C for 5 minutes and storing samples at -80 °C. Supernatants from these samples were used during western blot. The remaining bead suspension was placed on ice until the next step.

To purify RNA from immunoprecipitation samples, proteinase K buffer was prepped for the appropriate number of samples according to Magna RIP™ RNA-Binding Protein Immunoprecipitation Kit protocol. Beads from each immunoprecipitation samples were then resuspended in 150 µl of proteinase K buffer. RNA input samples were thawed and 107 µl of RIP wash buffer, 15 µl of 10% SDS, and 18 µl of proteinase K was added. All samples were incubated at 55 degrees for 30 minutes with gentle shaking to digest the protein. After the incubation, tubes were centrifuged briefly, and placed on the magnetic stand. Supernatants containing RNA were transferred into new tubes. To purify RNA, 250 µl of RIP wash buffer and 400 ul of acid phenol:chloroform:isoamyl alcohol (Thermo Scientific AM9720) was added to each tube. Tubes were vortexed at maximum setting for 15 seconds and centrifuged at 14000 rpm for 10 minutes at room temperature to separate the phases. 350 µl of the aqueous phase was removed, placed in a separate tube, and 400 µl of RNase free water added to the original tube. Tubes were then once again vortexed for 15 seconds and centrifuged at 14000 rpm for 10 minutes at room temperature to separate the phases. 300 µl of the aqueous phase was combined with the aqueous phase saved earlier. To precipitate RNA, 1/10 volume of 3M sodium acetate pH 5.1, 1.5 volume of isopropanol, and 1 µL of Glycoblue (Thermo Scientific AM9515) was added to each tube before mixing and incubating the tubes at -80 °C overnight. The next day, all tubes were centrifuged at 14000 rpm for 30 minutes at 4 °C and supernatant discarded. RNA pellet was washed once with 80% ethanol and tubes once again centrifuged at 14000 rpm for 15 minutes at 4 °C. Supernatants were then discarded carefully and pellets air dried. Pellets were resuspended in 15 ul of Rnase-free water, and RNA quantified using nanodrop. Equal amounts (in ng) of RNA were used for subsequent cDNA synthesis and qPCR (procedure described in Interferon-ß treatment time course experiments above). RNA samples were stored at -80 °C. To calculate percent input after qPCR analysis, the following formula was used: 100*2^(Adjusted input Ct – IP Ct). Adjusted input Ct was calculated using the following formula: Input Ct – log_2_(10) if 10% input was saved.

### Polysome profiling experiments

HEK293T cells were plated on 3×15-cm dishes at a density of 1E5 cells/cm^2^. The next day, one plate was transfected with 70 µg of DDX60 wt construct and two plates were transfected with 70 µg each of DDX60 E890A construct using TransIT®-LT1 Transfection Reagent at a 1:3 (µg to µl) DNA to lipid reagent ratio in a total volume of 3.5 mL of Opti-MEM according to the TransIT®-LT1 manufacturer’s protocol. 16-18 hours post transfection, DDX60 wt transfected cells were split 1 to 2 from 1×15-cm plate to 2×15-cm plate and DDX60 E890A transfected cells were split 1 to 2 from 2×15-cm plates to 4×15-cm plates. The next day, media on all the cells were changed to DMEM supplemented with 1.5% FBS, penicillin and streptomycin, and non-essential amino acids. Subsequently, one DDX60 wt transfected plate of cells and two DDX60 E890A transfected plate of cells were transfected with ∼30 – 40 pmol of 5’ cap-driven Fluc reporter mRNA and one DDX60 wt transfected plate of cells and two DDX60 E890A transfected plate of cells were transfected with ∼30 – 40 pmol of EMCV IRES-driven Fluc reporter mRNA. mRNA transfections were carried out using TransIT®-mRNA Transfection Kit at a 1:2 (µg to µl) RNA to lipid reagent ratio in a total volume of 3.5 mL of Opti-MEM according to the TransIT®-mRNA manufacturer’s protocol. The day after, one plate of DDX60 E890A and 5’ cap-driven Fluc reporter mRNA transfected cells and one plate of DDX60 E890A and EMCV IRES-driven Fluc reporter mRNA transfected cells were treated with 200 µM (∼0.1 µg/mL) of puromycin for 20 min at 37 °C by replacing the media on the cells with puromycin containing DMEM supplemented with 1.5% FBS, penicillin and streptomycin, and non-essential amino acids. Cells were then harvested. To harvest, all cells were first treated with 15 mL of complete growth media + cycloheximide (Sigma-Aldrich C7698) at 100 µg/mL for 15 min at 37 °C by replacing the media on the cells with cycloheximide containing DMEM supplemented with 10% FBS, penicillin and streptomycin, and non-essential amino acids. Cells were then washed once with 3 mL PBS + calcium and magnesium + cycloheximide at 100 µg/mL. Cells were then trypsinized with 3 mL trypsin + 100 µg/mL cycloheximide. When cells lifted off, 10 mL PBS + 100 µg/mL cycloheximide supplemented protease and phosphatase inhibitors (Thermo Scientific 87786 and Thermo Scientific 78420) were added and cells transferred to 15 mL tubes. Cells were pelleted by centrifugation at 1250 rpm for 4 min at 4 °C. Supernatant was aspirated and cells resuspended in 10 mL of PBS + cycloheximide supplemented protease and phosphatase inhibitors. 1 mL of cell suspension was collected in 1.5 mL tubes for input. Cells in both 15 mL and 1.5 mL tubes were pelleted by centrifugation at 1250 rpm for 4 min at 4 °C. Supernatants were again discarded, and all cell pellets frozen using liquid nitrogen before storing at -80 °C.

The day before running cell lysates through sucrose gradient, sucrose gradients were prepared by pipetting 5.5 mL of 50% sucrose solution (28.5 mL 1X LSB (20 mM Tris pH 7.4-7.5, 10 mM NaCl, 3 mM MgCl_2_), 21.5 g sucrose, 40 U/mL Riboblock (Thermo Scientific EO0382), 100 µg/mL cycloheximide) in a Beckman 14×89mm tube (Beckman Coulter 331372) and adding 5.5 mL of 15% sucrose solution (42.5 mL 1X LSB (20 mM Tris pH 7.4-7.5, 10 mM NaCl, 3 mM MgCl_2_), 8 g sucrose, 40 U/mL Riboblock, 100 µg/mL cycloheximide) on top. Enough gradients were prepared for the number of samples being processed. The tubes with the gradients were then sealed with parafilm and laid horizontally at 4 °C overnight. The next day, cell pellets were thawed on ice. Once pellets were thawed, one to two samples were processed at a time. 750 ul of 1X LSB + 100 µg/mL cycloheximide + 40 U/mL riboblock were added to the cell pellet and the pellet was incubated in a dounce homogenizer on ice for 3 min. Next, 250 µl of Triton X-100 detergent buffer (1.2% Triton X-100, 0.2 M sucrose (342 mg), 4.4 mL 1X LSB, 100 µg/mL cycloheximide, 40 U/mL riboblock) was added and cells were lysed by using a dounce homogenizer. Sample was then transferred to a 1.5 mL tube on ice. Once all pellets were lysed, samples were centrifuged at maximum speed for 5 minutes at 4 °C. The supernatants were then transferred to 1.5 mL tubes containing 100 µl of heparin solution (10 mg/mL heparin, 1.5 M NaCl, 1.4 mL 1X LSB, 100 µg/mL cycloheximide, 40 U/mL riboblock). RNA concentration in the tubes were then measured using a nanodrop. The sample volume being loaded into the sucrose gradients were removed from the gradients first and ∼1 mg of RNA was pipetted onto the gradients. Gradient tubes were balanced and ultracentrifuged at 36000 rpm for 2 hours at 4 °C using a SW 41 Ti rotor. Fractions from gradient tubes were then collected using a BR-188 Density Gradient Fractionation System (Brandel) into 1.5 mL tubes filled with 25 µl of 0.5M EDTA (Ambion AM9260G). All fractions were frozen at -80 °C.

RNA was isolated from ribosomal subunit and monosome fractions (1 - 3), low molecular weight (LMW) polysome fractions (4 and 5), and high molecular weight (HMW) polysome fractions (6 - 8) for each sample using an acid phenol chloroform extraction. Briefly, an equal volume of acid phenol chloroform was added, samples mixed 3-5 times by inverting tube up and down, and then vortexed to assure complete mix of phenol with sample. Samples were then centrifuge at maximum speed for 5 min at 4 °C. The upper aqueous phase was transferred to a new 2 mL tube. An equal volume of RNase free water was then added to the remaining phenol, samples mixed 3-5 times by inverting tube up and down, vortexed, and centrifuge at maximum speed for 5 min at 4°C. The aqueous phase from this spin was combined with the previously collected aqueous phase. To precipitate RNA, 1/10 volume of 3M sodium acetate pH 5.1, 1.5 volume of isopropanol, and 1 µL of Glycoblue was added to each tube before mixing and incubating the tubes at -80 °C overnight. The next day, all tubes were centrifuged at maximum speed for 30 minutes at 4 °C and supernatant discarded. RNA pellet was washed twice with 1 mL of 70% ethanol, centrifuging at maximum speed for 15 minutes at 4 °C in between washes. Supernatants were then discarded and pellets air dried. Pellets were resuspended in 20 ul of Rnase-free water. An equal volume of RNA from each fraction was used for cDNA synthesis (procedure described in Interferon-ß treatment time course experiments above) and qPCR analysis was carried out as described in (Panda et al., 2017).

### Quantification and statistical analysis

All replicates of *in vitro* experiments are from biologically independent experiments unless directly stated. Statistical analyses were performed in Prism (GraphPad Prism Version 9.3.1 (350). The statistical tests used, and number of biological replicates are indicated in each figure legend. Statistical significance was defined as a p value of 0.05.

## Author contributions

### Conceptualization

Mohammad Sadic, William M. Schneider, Olga Katsara, Gisselle N. Medina, Aishwarya Mogulothu, Yingpu Yu, Meigang Gu, Teresa de los Santos, Robert J. Schneider, Meike Dittmann

### Data curation

Mohammad Sadic, Olga Katsara, Gisselle N. Medina, Aishwarya Mogulothu, Meike Dittmann

### Formal analysis

Mohammad Sadic, Olga Katsara, Giselle N. Medina, Aishwarya Mogulothu, Meike Dittmann

### Funding acquisition

Teresa de los Santos, Robert J. Schneider, Meike Dittmann

### Methodology

Mohammad Sadic, William M. Schneider, Olga Katsara, Gisselle N. Medina, Aishwarya Mogulothu, Yingpu Yu, Meigang Gu, Teresa de los Santos, Robert J. Schneider, Meike Dittmann

### Supervision

Teresa de los Santos, Robert J. Schneider, Meike Dittmann Writing – original draft: Mohammad Sadic, Robert J. Schneider, Meike Dittmann

### Writing – review&editing

Mohammad Sadic, William M. Schneider, Olga Katsara, Gisselle N. Medina, Aishwarya Mogulothu, Yingpu Yu, Meigang Gu, Teresa de los Santos, Robert J. Schneider, Meike Dittmann

## Acknowledgments

We would like to thank Charles M. Rice for mentorship and support to initiate this project, bicistronic and monocistronic HCV reporters, and constructs used in (Schoggins et al., 2011) screen; Vincent Racaniello and Eckard Wimmer for strategizing the cloning and generation of chimeric poliovirus (EMCV-IRES-PV); Ann Palmenberg for pEC9 plasmid and pPVM-2A-144-poliovirus-GFP plasmid; Aaron Briley, Adil Mohamed, Maren de Vries, Ana M. Valero-Jimenez, Keaton Crosse, Rachel Prescott, Austin Schinlever, and Ralf Duerr for helpful discussions; Aaron Briley, Sarah Ballentine, Chloe Adrienna Talana, and Paige Loose for technical assistance.

